# *In silico* evolution of protein binders with deep learning models for structure prediction and sequence design

**DOI:** 10.1101/2023.05.03.539278

**Authors:** Odessa J Goudy, Amrita Nallathambi, Tomoaki Kinjo, Nicholas Randolph, Brian Kuhlman

## Abstract

There has been considerable progress in the development of computational methods for designing protein-protein interactions, but engineering high-affinity binders without extensive screening and maturation remains challenging. Here, we test a protein design pipeline that uses iterative rounds of deep learning (DL)-based structure prediction (AlphaFold2) and sequence optimization (ProteinMPNN) to design autoinhibitory domains (AiDs) for a PD-L1 antagonist. Inspired by recent advances in therapeutic design, we sought to create autoinhibited (or masked) forms of the antagonist that can be conditionally activated by proteases. Twenty-three *de novo* designed AiDs, varying in length and topology, were fused to the antagonist with a protease sensitive linker, and binding to PD-L1 was tested with and without protease treatment. Nine of the fusion proteins demonstrated conditional binding to PD-L1 and the top performing AiDs were selected for further characterization as single domain proteins. Without any experimental affinity maturation, four of the AiDs bind to the PD-L1 antagonist with equilibrium dissociation constants (K_D_s) below 150 nM, with the lowest K_D_ equal to 0.9 nM. Our study demonstrates that DL-based protein modeling can be used to rapidly generate high affinity protein binders.

**Significance statement:** Protein-protein interactions are critical to most processes in biology, and improved methods for designing protein binders will enable the creation of new research reagents, diagnostics, and therapeutics. In this study, we show that a deep learning-based method for protein design can create high-affinity protein binders without the need for extensive screening or affinity maturation.

## Introduction

Engineered proteins that bind to proteins of interest are useful as research reagents, diagnostics, and therapeutics. Over the last 15 years, computational protein design has emerged as an effective approach for designing protein binders(1–4). Until very recently, these methods have been based on atomistic models of proteins in which the relative favorability of different sequences and conformations is evaluated with energy functions that model physical phenomena such as van der Waals forces and hydrogen bonding(5). One attractive feature of these methods is that they allow the binding site to be specified on the target protein. However, following the traditional *in silico* design process, it is frequently necessary to screen hundreds or thousands of designed proteins to identify binders, and then use experimental affinity maturation to optimize the sequences to bind with equilibrium dissociation constants (K_D_s) below 100 nM(1, 4, 6).

In the last few years, advances in deep learning (DL) have dramatically improved computational methods for protein modeling. Structure prediction networks such as AlphaFold2 (AF2)(7) and RoseTTAFold(8) have been trained to accurately predict protein structure from sequence information, while design networks such as ProteinMPNN(9) identify amino acid sequences that are compatible with a given protein backbone(10–12).

Here, we introduce a protein design pipeline called EvoPro that uses a genetic algorithm comprised of iterative rounds of structure prediction with AF2 and sequence diversification to evolve a set of proteins to bind a pre-specified target protein (Fig. 1A). First, a set of *de novo* designed miniprotein sequences previously engineered to fold into well-defined structural motifs(1) are passed to AlphaFold-Multimer(13) (a version of AF2 trained to predict the structures of protein-protein complexes) along with the target protein sequence. Next, to identify the more favorable sequences, the predicted AF2 complexed structures are evaluated with a fitness function derived from AF2 confidence scores and the number of interface contacts between the designed binder and the target protein (Fig. 1A, right). The top-scoring sequences are diversified for the next generation using either random mutagenesis and crossover or are optimized with ProteinMPNN using the predicted structures of the complex as input (Fig. 1A, bottom). With each generation, the fitness function selects for sequences with corresponding predicted structures that have higher AF2 confidence scores and better interface interactions at the desired surface patch on the target protein.

**Figure 1.**
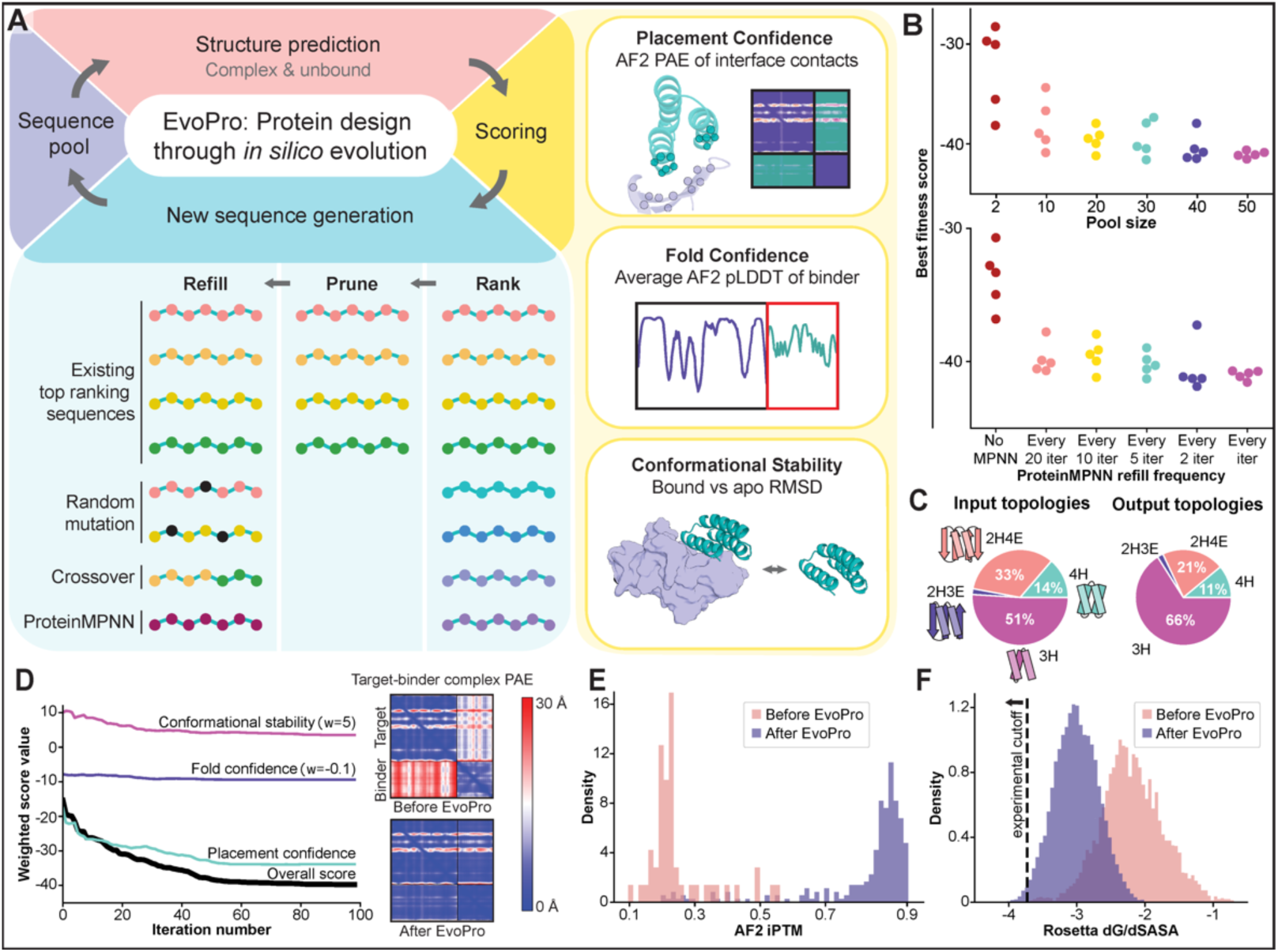
EvoPro design pipeline. **(A)** EvoPro is a cyclic design process that iterates between sequence generation and scoring through structure prediction within a genetic algorithm framework. Sequence diversification is done through random mutagenesis or ProteinMPNN (bottom, in blue), while structure prediction is performed with AF2. The scoring step includes three terms calculated from AF2 predictions (right, in yellow): Placement confidence score, which uses AF2 PAE confidences to estimate the quality of the interface; Fold confidence score, which uses AF2 pLDDT confidences to estimate the miniprotein’s folding stability; and Conformational stability score, which minimizes the conformational difference between bound and unbound forms of the protein binder. **(B)** Increasing the size of the sequence pool and the frequency of sequence optimization with ProteinMPNN produces more consistent and lower overall fitness scores. Each dot represents an independent design trajectory. **(C)** All of the topologies represented in the 51 starting scaffolds are present in the top 100 output models as ranked with the AF2 ipTM score (a global metric of interface confidence). **(D)** Score terms as a function of iteration number for a single trajectory of EvoPro. Weight on score term indicated with w = x. On the right, PAE heatmaps from AF2 predictions of the target-binder complex before and after EvoPro optimization. Lower PAE indicates more confidence. **(E)** Distribution of AF2 ipTM scores before and after EvoPro optimization. A larger score indicates higher confidence. **(F)** Change in Rosetta dG/dSASA, which is an independent metric of interface quality, after EvoPro optimization. Lower energies are more favorable.

An attractive feature of the EvoPro pipeline is that, through iterative structure prediction and sequence design, scaffolds can undergo conformational changes that may be favorable for binding. Coupling backbone plasticity with interface design has previously been challenging to encode in more traditional computational design methods. EvoPro is conceptually similar to other design pipelines that have been recently described, such as hallucination with RoseTTAFold(14) and AlphaDesign(15) that combines AF2 with random mutagenesis to design proteins. One component that differentiates EvoPro from these previous studies is the iterative exchange between ProteinMPNN and AF2. Sequences created with AlphaDesign have not been experimentally validated, while RoseTTAFold-based hallucination has been used to create new protein structures, assemblies, and proteins that bind tightly to helical peptides(14, 16, 17).

To demonstrate EvoPro’s utility, we designed various autoinhibitory domains (AiDs) for an antagonist of PD-L1, a critical component of the programmed death pathway and a clinically relevant immunotherapeutic target. The binding of PD-1 on T cells to PD-L1 on adjacent cells downregulates T-cell inflammatory activity, and cancer cells frequently overexpress PD-L1 to suppress the anti-tumor immune response(18). Monoclonal antibodies that bind to either PD-1 or PD-L1 and block the endogenous interaction are successful treatments against several types of cancer but are also associated with toxic side effects, as T-cell activity is systemically enhanced outside the tumor microenvironment(19). One promising strategy that is being tested in clinical trials for mitigating the toxic effects of PD-1/PD-L1 inhibitors is engineering autoinhibitory (or masking) domains (AiDs) that block the activity of the drug until it reaches the tumor microenvironment(20–22). Using this approach, the masking domain blocks the binding loops of the therapeutic antibodies and weakens affinity for the target until a linker is cleaved by tumor-enriched proteases(23, 24).

In this work, instead of using an antibody as a PD-L1 antagonist, we employ a soluble variant of PD-1 that has been affinity matured to bind tightly to PD-L1 (K_D_ < 1 nM)(25, 26). This antagonist, named HA-PD1 for <u>h</u>igh <u>a</u>ffinity <u>PD</u>-<u>1</u>, has been shown to shrink tumors in animal models(25). To regulate the activity of HA-PD1 so that it has weak affinity for PD-L1 until activated with proteases, we used EvoPro to design a set of diverse small miniprotein domains as AiDs that can be fused to HA-PD1 and block its interaction with PD-L1. Binding to PD-L1 is restored when the linkers connecting HA-PD1 to the AiDs are cleaved with protease treatment. Moreover, when expressed as separate domains, several of the recombinant AiDs bind to HA-PD1 with K_D_ values lower than 150 nM. Overall, these results demonstrate that EvoPro is an effective method for designing high affinity protein binders.

## Results

### EvoPro architecture

The EvoPro pipeline uses iterative rounds of structure prediction with AF2 and sequence design with ProteinMPNN to identify favorable regions of sequence-structure space. We use a runtime-optimized protocol for AF2, which allows structure predictions in 5-10 seconds rather than minutes (see Methods). As ProteinMPNN runs in just ∼1 second per generated sequence, we do not perform any runtime optimization for use in a high-throughput manner. Therefore, during each EvoPro iteration, we can predict structures for many sequences within a few minutes.

Each EvoPro trajectory attempts to meet predefined design requirements by selecting and evolving sequences with the best (i.e., lower) fitness scores (Fig. 1A). To use EvoPro for binder design, we incorporated score components representing interface quality (“Placement Confidence”), binder folding stability (“Fold Confidence”), and binder conformational difference of the bound versus unbound states (“Conformational Stability”) (Fig. 1A, right). Notably, alternative score terms can be implemented and combined for other design problems, and we are currently exploring this further.

We performed benchmarking simulations with various pool sizes (i.e., number of amino acid sequences in the population) and schemes for sampling sequence space (Fig. 1B). Trajectories with smaller pool sizes were unable to consistently produce good designs with desirable fitness scores, frequently getting stuck in local fitness minima. Increasing the pool size generally results in lower fitness scores, but also comes with a higher computational expense as each sequence must have its structure predicted.

In each generation, EvoPro introduces mutations into the best-scoring sequences through random mutagenesis, crossover, or optimization with ProteinMPNN (Fig. 1A, bottom). When using ProteinMPNN, the AF2-predicted model of the binder:HA-PD1 complex served as the input, and mutations were allowed at all residue positions in the binder. Incorporating ProteinMPNN in the pool refill step significantly decreases the overall minimum fitness score achieved per trajectory (Fig. 1B, bottom). For the design of the PD-L1 antagonist autoinhibitory domains, we used ProteinMPNN once every 10 iterations to refill the pool. However, later benchmarking showed that more frequent use of ProteinMPNN often leads to even lower overall fitness scores.

### Generating autoinhibitory domains with EvoPro

To design autoinhibitory domains for the PD-L1 antagonist, we selected a set of 51 topologically diverse miniprotein scaffolds (Table S1) from the recommended set of *de novo* designed miniprotein scaffolds described by Cao *et al.*^1^. For each of these 51 starting scaffolds, we ran five independent EvoPro design trajectories, each with a pool size of 50 sequences over 60 iterations. In total, 255 design trajectories were performed with each trajectory generating a final pool of 25 designed sequences, leading to 6,375 total designs. Each independent EvoPro trajectory tends to converge to a set of sequences that are highly similar, with an average sequence identity of 81%, representing a local minimum in fitness space.

Four diverse scaffold topologies were used as starting points for EvoPro trajectories, including three-helix (3H) and four-helix bundles (4H) as well as miniproteins with the mixed topologies of two helices with three β-strands (2H3E) and two helices with four β-strands (2H4E) (Fig. 1C, Table S1). After EvoPro optimization, the top 100 results (as ranked by AF2’s interface predicted template modeling score (ipTM)) include each topology, although the proportion of 3H bundles is slightly greater (Fig. 1C). This may indicate that either this topology is particularly well suited to the target binding interface or that there is some bias towards this topology in the DL models that were used.

A representative EvoPro design trajectory shows the individual score components decreasing as the simulation progresses over iterations (Fig. 1D). The three score components are: 1) the placement confidence score, which represents the size and quality of the interface as derived from AF2’s pairwise predicted aligned error (PAE); 2) the fold confidence score, which is based on AF2’s residue-based confidence scores (pLDDT) of the miniprotein binder; and 3) the conformational stability score, which represents the conformational difference between the binder in its monomeric form versus complex form and minimizes the conformational change required for binding (see Methods) (Fig. 1A, right).

The placement confidence score, which represents the quality of the interface, shows the most drastic improvement over the simulation, indicating that the initially poor interfaces evolved into better ones (Fig. 1D, left). AF2 PAE heatmaps before and after EvoPro illustrate improved confidence in the designed interface (Fig. 1D, right). The conformational stability score gradually decreases, favoring sequences predicted to have minimal structural changes upon binding (Fig. 1D, left). The fold confidence score shows the least change over the trajectory, likely because the scaffolds that were selected are known to be quite stable and, therefore, have high per-residue AF2 confidences (pLDDT) at the start of the trajectory (Fig. 1D, left). The AF2 ipTM confidence metric, which represents global interface confidence and was not optimized during the design simulations, also shows a drastic improvement for target-binder interfaces after EvoPro optimization (Fig. 1E).

We orthogonally verified the interface quality of our designs using Rosetta-based scoring metrics. Following FastRelax(27) rotamer optimization and backbone minimization, the InterfaceAnalyzer mover(28) was applied to calculate interface energy between the binder and target protein. The interface quality metric dG_separated/dSASA × 100 (dG/dSASA) was used to score the 6,375 designed AiDs. The score distribution shifts towards lower (i.e., better) energy interfaces after EvoPro optimization, as compared to the interfaces of the starting miniprotein scaffolds (Fig. 1F). This metric was then used to sort and select the 100 best designs. For these, the resulting models from AF2/EvoPro were then compared to predictions made with another DL-based method, OmegaFold(29). These predictions were made with a 28-amino acid flexible linker connecting the C-terminus of HA-PD1 to the N-terminus of the miniprotein to resemble the final autoinhibited complexes we aimed to build. Sequences with similar AF2 and OmegaFold models (as determined by RMSD < 3Å) were further filtered to reduce redundancy of their starting scaffolds to yield a final set of 23 designs for experimental characterization (Table S2). The final TEV protease-cleavable linker for each construct was optimized using OmegaFold to generate the 23 masked antagonist (MA) constructs (Table S3, see Methods).

Despite binding to the same surface patch on HA-PD1, the specific contacts made between each AiD and HA-PD1 differ among the designs and from the HA--PD1:PD-L1 interactions (Fig. 2). This indicates that the design process is not simply recapitulating the native contacts but is instead creating novel *de novo* interfaces. However, the major “rules” of the interface are still fulfilled by the diverse models: a hydrophobic patch on HA-PD1 is complemented by hydrophobic residues on the binder, and polar contacts are made around the periphery of the interface, including intermolecular hydrogen bonds involving H64_HA-PD1_, H68_HA-PD1_, and D85_HA-PD1_. Additionally, there is a pocket in the center of the HA-PD1 binding surface (Fig. S1) that is filled by Y123_PD-L1_. Many of the designed binders include a tyrosine or leucine that is inserted in the same pocket (Fig. S1).

**Figure 2.**
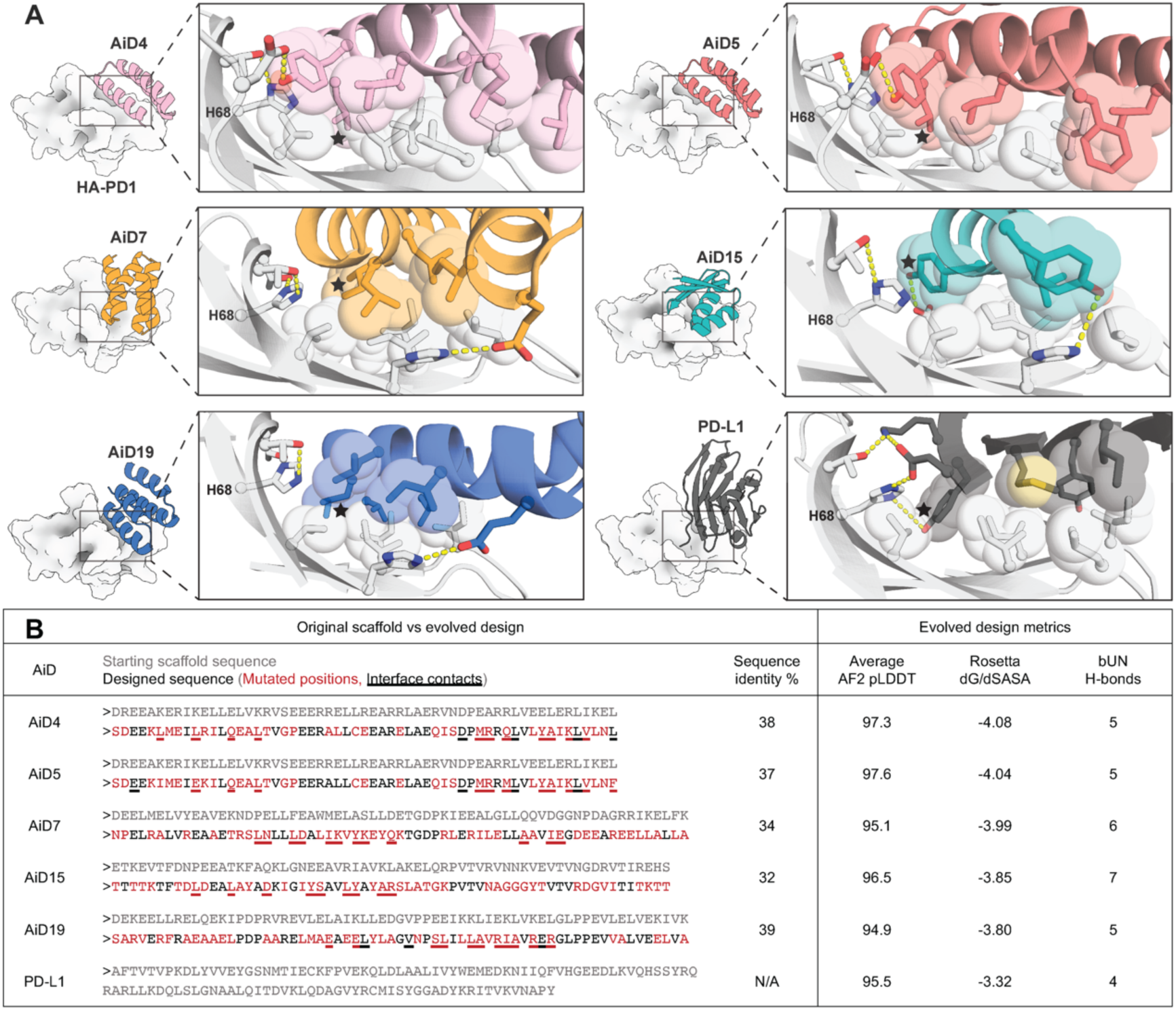
AF2 models of designed interface contacts are different than those at the HA-PD1:PD-L1 interface. **(A)** sharing a similar surface patch on HA-PD1, the designs make unique and different contacts than that of the natural ligand, PD-L1 (PDB 5IUS). One common feature, however, is the insertion of a hydrophobic residue (marked with a black star) into a hydrophobic pocket in HA-PD1. The AF2 models were energy minimized with Rosetta before generating the figures. Figure made in PyMOL where some residues were hidden for clarity and H68_HA-PD1_ is shown in every panel. **(B)** The table compares the designed sequences with the corresponding starting scaffold sequence provided as input to EvoPro. Scoring metrics for the design models are also provided for the Rosetta dG/dSASA and buried unsatisfied hydrogen bonds (bUN H-bonds) parameters.

### Masked antagonists display protease-dependent binding to PD-L1

The twenty-three MAs were expressed at a small scale using Expi293 mammalian cells. Secreted proteins were purified using nickel resin and the eluates were evaluated for yield and purity (Fig. S2). Proteins of the appropriate molecular weight were observed for all designs, but expression yields were over 10-fold greater for some constructs. Thirteen well-expressing constructs that represented a variety of scaffold topologies were selected for additional screening (Fig. S2) using a cell-based PD-L1 binding assay developed by Promega. The assay detects interactions between PD-1 displayed on engineered Jurkat T-cells and PD-L1 displayed on engineered CHO cells. If the interaction between the endogenously expressed PD-1 and PD-L1 is disrupted, T-cell receptor (TCR) signaling is activated and drives NFAT-mediated luciferase activity. The PD-L1 blockade assay was performed for the 13 masked antagonists at two concentrations (10 and 100 nM) in the presence and absence of TEV protease (Fig. S3, S4A). Eight of the masked antagonists (MA4, MA7, MA9, MA10, MA15, MA17, MA19, and MA20) showed conditional PD-L1 binding upon protease treatment (Fig. S3). Moreover, MA5 uniquely displayed reduced binding activity both with and without protease treatment (Fig. S3), suggesting that the AiD did not dissociate following protease treatment, due to particularly tight affinity for HA-PD1.

To further characterize the top candidates (MA4, MA5, MA7, MA9, MA10, MA15, MA16, MA17, MA19, and MA20), they were expressed at a larger scale and purified using nickel affinity and size exclusion chromatography (Fig. S5). As MA16 did not show any masking in the preliminary experiments, it was chosen as a negative control. For all designs, the final yields were favorable (an average of ∼60 µg protein/mL of cell medium). Three of the designs (MA4, MA5, and MA15) eluted as two peaks from the gel filtration column (Fig. S5), perhaps indicating domain swapping where the AiD from one chain interacts with HA-PD1 from another chain. Protein from the second peak (longer retention time) was used in all experiments.

The Promega cell-based PD-L1 binding assay was repeated but performed at a wider concentration range to determine IC50 values with and without protease treatment (Fig. 3A, S4B). HA-PD1 without a mask (NoMA) inhibited the endogenous cell surface interaction between PD-1 and PD-L1 with an IC50 of 16 nM, comparable to checkpoint inhibitors currently used in the clinic. Prior to protease treatment, several of the masked antagonists (MA5, MA10, MA15, MA19, and MA20) were poor inhibitors with IC50s near or above 1 µM, indicating that their respective AiDs prevent HA-PD1:PD-L1 binding as designed (Fig. 3B, S6, Table 1). Pretreatment with protease rescued activity for many of the inhibitors, and the IC50s lowered to values below 30 nM. As observed in the preliminary screen, MA5 was a poor inhibitor even after protease treatment (IC50 = 582 nM) (Fig. 3B, Table 1), consistent with the AiD remaining bound to HA-PD1 after cleavage of the linker.

**Figure 3.**
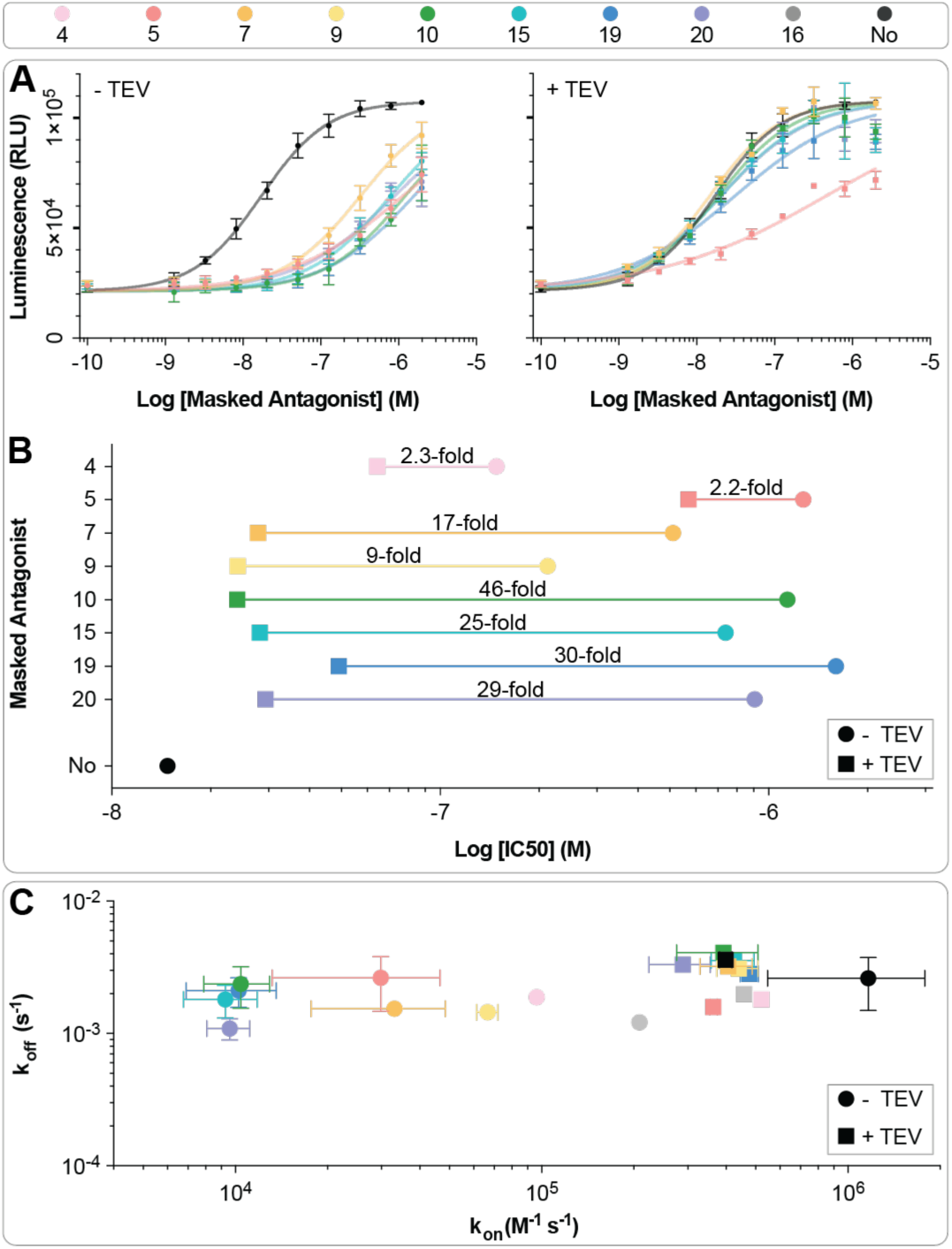
EvoPro binders serve as autoinhibitory domains for a PD-L1 antagonist. In the cell assay, competitive inhibition of the native PD-1:PD-L1 interaction drives luciferase expression, which is detected by monitoring luminescence. **(A)** The competitive binding of various masked PD-L1 antagonists (as indicated in the color key) were measured as a function of protein concentration with (right) and without (left) protease treatment. The activity of HA-PD1 without a mask (“No”) is shown for reference in black. Two technical replicates from a single experiment, presented as mean ± SEM are shown. **(B)** IC50s of the masked antagonists from the cell surface assay with (squares) and without (circles) protease treatment. Additional data from two independent experiments are included in the Supporting Information (Fig. S6). **(C)** Association rate constants (k_on_) and dissociation rate constants (k_off_) for the masked antagonists binding to PD-L1 with (squares) and without (circles) protease treatment. Binding measurements were made with SPR with biotinylated PD-L1 immobilized on a NeutrAvidin chip (Fig. S7). Reported error represents SD from at least 3 independent experiments except for MA4 (n = 1), MA16 (n = 1), and MA20 (n = 2).

**Table 1.** Masked antagonists show protease-dependent binding and activity. Table of mean kinetic parameters for the masked antagonists before and after protease treatment including NoMA (“No”) for the SPR binding kinetic assay (left) and the cellular activity assay (right). For the binding kinetics, error represents SD whereas some samples were only tested once (no replicate, n.r.). For the cell assay, error represents either the SD in the average IC50 across multiple experiments (Fig. S6) or the 95% confidence interval of 3 replicates within a single experiment (*).

| MA | SPR |  |  | Cell Assay |
| --- | --- | --- | --- | --- |
| | $k_{on} (M^{-1} s^{-1}) / 10^3$ | $k_{off} (s^{-1}) \times 10^5$ | $K_D$ (nM) | IC50 (nM) |
| 4 | 96.3 (n.r.) | 189 (n.r.) | 19.7 (n.r.) | 149 ± 17.8* |
| 4 + TEV | 522 (n.r.) | 181 (n.r.) | 3.5 (n.r.) | 64 ± 10.7* |
| 5 | 30 ± 16.7 | 264 ± 116 | 122 ± 91 | 1285 ± 214 |
| 5 + TEV | 364 ± 10.4 | 159 ± 7.6 | 4.4 ± 0.3 | 582 ± 171 |
| 7 | 33.1 ± 15.4 | 155 ± 8.5 | 61.4 ± 43.3 | 566 ± 342 |
| 7 + TEV | 408 ± 78.7 | 321 ± 6 | 8.1 ± 1.8 | 34 ± 26.7 |
| 9 | 66.6 ± 5.3 | 145 ± 11 | 21.8 ± 0.5 | 212 ± 10.4* |
| 9 + TEV | 441 ± 67.9 | 311 ± 9.6 | 7.2 ± 1 | 24 ± 4.8* |
| 10 | 10.4 ± 2.5 | 238 ± 82.6 | 243 ± 113 | 1143 ± 102 |
| 10 + TEV | 392 ± 117 | 410 ± 26.7 | 11.1 ± 3.2 | 25 ± 8.5 |
| 15 | 9.3 ± 2.5 | 181 ± 50.1 | 205 ± 68.3 | 743 ± 68.7 |
| 15 + TEV | 425 ± 67.7 | 355 ± 8.2 | 8.5 ± 1.2 | 30 ± 12.8 |
| 16 | 209 (n.r.) | 122 (n.r.) | 5.8 (n.r.) | N/A |
| 16 + TEV | 459 (n.r.) | 199 (n.r.) | 4.3 (n.r.) | N/A |
| 19 | 10.3 ± 3.3 | 211 ± 52.8 | 228 ± 102 | 1606 ± 45.3 |
| 19 + TEV | 478 ± 32.3 | 283 ± 4.4 | 5.9 ± 0.5 | 54 ± 32.9 |
| 20 | 9.6 ± 1.5 | 109 ± 20.3 | 114 ± 13.8 | 909 ± 33 |
| 20 + TEV | 290 ± 65.8 | 332 ± 7.8 | 11.7 ± 2.4 | 31 ± 15.7 |
| No | 1165 ± 617 | 262 ± 113 | 2.5 ± 1 | 16 ± 6.6 |
| No + TEV | 400 ± 18.2 | 363 ± 29.3 | 9.1 ± 0.9 | N/A |

The equilibrium dissociation constants (K_D_s) of the MAs for PD-L1 were measured with surface plasmon resonance (SPR) (Fig. 3C, Table 1) before and after incubation with TEV protease (Fig. S7). Biotinylated PD-L1 was immobilized on the sensor chip and single-cycle kinetics were used to monitor binding as a function of the concentration of the MA. Prior to protease cleavage, many MAs displayed reduced affinity for PD-L1. Uncleaved MA5, MA10, MA15, MA19, and MA20 all bound PD-L1 with K_D_ values near 100 nM while NoMA bound with a K_D_ of 2.5 nM. Following protease treatment, affinity for PD-L1 was restored in almost all cases and the measured K_D_s were below 10 nM (Fig. 3C, Table 1). Unlike in the cell-based assay, protease-treated MA5 bound to PD-L1 with an affinity similar to NoMA, which is consistent given the experimental setup of each technique. During the SPR experiments, the buffer is flowing continually over immobilized PD-L1 and, therefore, during the dissociation phase, the cleaved AiD is removed, preventing it from rebinding to HA-PD1 and blocking binding to PD-L1. In the cell-based assay, the cleaved AiD remains in solution and can rebind to HA-PD1, blocking binding to PD-L1.

The SPR data were also used to determine the association and dissociation rate constants (k_on_ and k_off_, respectively) of the masked antagonists for PD-L1. Interestingly, all the constructs, regardless of whether they had been treated with protease, dissociated from PD-L1 at a similar rate that HA-PD1 dissociates from PD-L1 (Fig. 3C, S7, Table 1). As the original HA-PD1 domain remained unmodified within the MA complex, binding to PD-L1 would be similar across all constructs. In contrast, the k_on_ values differ between constructs and reflect the extent each AiD prevents binding to PD-L1. MA5, MA10, MA15, MA19, and MA20 have k_on_ values < 30 × 10^3^ M^-1^s^-1,^ while NoMA has a k_on_ value > 1000 × 10^3^ M^-1^s^-1^. MA16, which did not display autoinhibition in the cell binding assay, also did not show reduced affinity for PD-L1 in the SPR experiments.

### Characterization of the AiDs as separately expressed proteins

To directly measure the affinity of the AiDs for HA-PD1, we selected a subset of AiDs to express as isolated domains: AiD4, AiD5, AiD7, AiD9, AiD10, AiD15, AiD19, and AiD20. All constructs, except for AiD20, expressed well and were purified using nickel affinity chromatography (Fig. S8A). All the constructs designed to be helical bundles have circular dichroism (CD) spectra consistent with the formation of α-helices with minima at 222 nm and 208 nm (Fig. 4A, S9). AiD15, which is designed to adopt an α/ý fold, has a minimum at 215 nm which is characteristic of β-strands. Like the parent sequences,(1) the AiDs demonstrated high thermostability and all of them except for AiD19 and AiD9 remained folded up to 90°C (Fig. 4B, S9).

**Figure 4.**
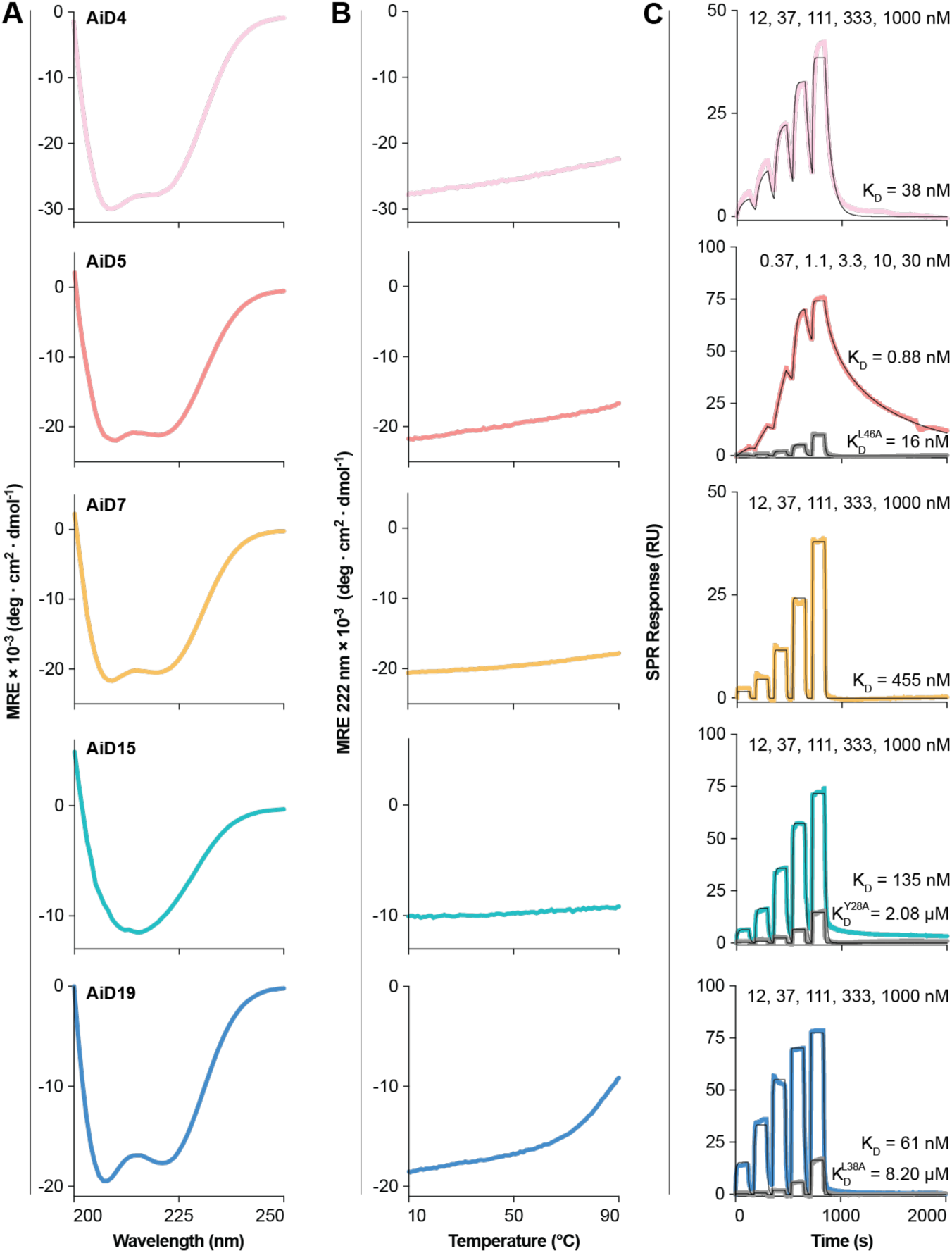
AiDs expressed as single domain proteins are folded and bind to HA-PD1. **(A)** Circular dichroism (CD) spectra and **(B)** temperature melts of various AiDs. **(C)** Single-cycle SPR sensorgrams for designed AiDs (colored line) and select mutant (grey line). Biotinylated HA-PD1 was immobilized on a NeutrAvidin chip to a level of ∼250 response units (RUs). The AiDs and mutants were injected at the indicated concentrations and the data were fit to a 1:1 binding model (black). The indicated K_D_ values are the average from three or more binding measurements (Table S4, Fig. S10, S12).

The binding affinities between the AiDs and HA-PD1 were measured using SPR, with HA-PD1 biotinylated and immobilized on the sensor chips. AiD5, which inhibits HA-PD1 in the cell binding experiment even after protease treatment, has the tightest affinity for HA-PD1 with a K_D_ of 0.88 nM (Fig. 4C, S10, Table S4). AiD7, which is a less potent mask in the cell binding assay, has a K_D_ of 460 nM. AiD4, AiD15, and AiD19 have K_D_ values of 38, 135, and 61 nM, respectively (Fig. 4C, Table S4). For all the AiDs, the rates of association were high with k_on_ > 1 × 10^5^ M^-1^s^-1^. The rates of dissociation were also rapid for many of the binders, with k_off_ > 0.1 s^-1^ for AiD7, AiD15, and AiD19 (Table S4, Fig. S10). We were unable to achieve consistent results for AiD9 and AiD10 in the SPR experiments.

To validate the AF2 models of AiD5, AiD15, and AiD19 bound to HA-PD1, we mutated select residues in the AiDs (Fig. S11) to alanine and measured binding affinities with SPR (Fig. 4C, S8B, Table S4). In all three cases, mutating the residue predicted to pack into the pocket occupied by Y123 in the PD-L1/HA-PD1 complex (AiD5:L46A, AiD15:Y28A, and AiD19:L38A) led to more than a 10-fold increase in K_D_ (Fig. 4C). Visual inspection was used to select an additional interface residue to mutate for each AiD. All three of the other single mutation variants (AiD5:Y49A, AiD15:L27A, and AiD19:L29A) (Fig. S11) increased K_D_ by more than 10-fold (Fig. S12). Moreover, since AiD5 binds exceptionally tight to HA-PD1, we selected two additional interface residues from AiD5 to mutate (Fig. S11). The single mutations, E10A and Q14A, increased the K_D_ from 0.9 nM to 26 nM and 4.4 nM respectively (Table S4, Fig. S12). These residues are predicted to be at the edge of the interface and form hydrogen bonds with backbone atoms on HA-PD1 (Fig. S11).

## Discussion

Our results demonstrate that DL-models for structure prediction and sequence design can be combined to design high-affinity protein binders without the need for experimental affinity maturation. Notably, the AiDs bind to HA-PD1 with a wide range of binding affinities. AF2 confidence scores and binding energies calculated with Rosetta were not able to predict that AiD5 would bind HA-PD1 more than 40-fold tighter than the other designs. It is interesting to compare AiD5 and AiD4 because the two sequences were produced by the same EvoPro trajectory and have only four amino acid differences at the interface (Fig. 2). This result highlights the continued need for improved methods for calculating protein-protein binding energies.

One powerful feature of EvoPro is that the score function can be customized to reward sequences that meet predefined requirements. In this study, we included a score term that favors binder sequences predicted to form the same structure in the unbound state as in the bound state. We included this score term, the conformational stability score, because changes in conformation upon binding come with an energetic penalty. A notable feature of the AiDs is that they all have rapid on rates with k_on_ values greater than 1 × 10^5^ M^-1^ s^-1^. This result is consistent with an interaction that does not involve large conformational changes as slower on rates are frequently observed for systems that undergo structural rearrangement upon binding(30). Without the score term that compares the unbound state to the bound state, EvoPro frequently produced sequences predicted to change conformation upon binding. This behavior may be enhanced by rewarding it in the score function and may be an exciting approach for designing allostery.

In designing AiDs for HA-PD1, EvoPro benefited from AF2’s tendency to dock proteins against the PD-L1 binding site on HA-PD1. This may reflect a desire to bury hydrophobic surface area on HA-PD1 as well as memory of the natural PD-1/PD-L1 binding site, since the AF2 training set most likely included the structure of PD-1 bound to PD-L1. In preliminary simulations with other target proteins, we have observed that EvoPro readily finds binders that interact with known sites of protein-protein interactions but struggles to place binders against other regions on the surface of target proteins (even if the score function rewards the alternative binding sites). This feature of EvoPro is favorable if the goal is to create competitive inhibitors that compete with a naturally occurring interaction but may be a problem when a novel binding surface is targeted. One exciting feature of RFDiffusion(31), an alternative DL-based approach that was recently applied to the design of protein binders, is that it enables directed docking at specified locations on a target protein.

In the cell-based assays, we observed that the AiD with the tightest affinity for HA-PD1 (AiD5) was not the most effective switch as the AiD remained bound to HA-PD1 following protease cleavage of the linker. Previously, we observed a similar result when using a soluble variant of PD-L1 as an AiD for HA-PD1(32). Masked PD-L1 antagonists currently being tested in humans show changes in affinity for PD-L1 in a range similar to MA15 and MA19(21). These findings highlight the utility of generating a panel of binders with varied affinity when building autoinhibited systems as the optimal switch may depend on the biological application.

## Materials and Methods

### Genetic algorithm

The EvoPro pipeline incorporates a method of solving optimization problems called a genetic algorithm that is modeled off the theory of natural selection and biological evolution(33). Genetic algorithms involve a population – in this case a pool of design sequences that could fold into a functional protein – being tested for fitness and evolving towards a more “fit” population. For our purpose, fitness signifies how well the designed sequence can fold into a functional protein of interest. Reminiscent of the concept of survival of the fittest, the algorithm evaluates the fitness of each sequence in a population based on predefined metrics and ranks the population from highest fitness to lowest. The lower half of the pool – the least fit – are removed and replaced with new “evolved” sequences as input to the next iteration of the algorithm.

### AlphaFold2 (AF2)

We use a local installation of AF2 which is optimized to run several sequences in parallel on multiple GPUs using a distribution script. The out-of-the-box runtime for a single sequence prediction through AF2 is on the order of tens of minutes, even on GPUs and using MMseqs2-based multiple sequence alignment (MSA) and template generation(34). To reduce this runtime to make high-throughput structure prediction more feasible, we remove the MSA calculation step by providing a pre-computed MSA for the target protein, which does not change throughout the design simulation, and use an empty MSA for the miniprotein scaffold. Previous studies have demonstrated that structure prediction models such as AF2 or RoseTTAFold can accurately predict the structure of small *de novo* proteins without an MSA(14). Removal of the MSA generation step brings the time per prediction down to ∼5 minutes. Within this, most of the time is spent in the compilation step of the AF2 model. Therefore, we incorporate a way to compile AF2 only once and keep it compiled on GPUs, waiting for the next sequence to be passed through. After the single compilation step, which takes ∼300 seconds, passing one multimeric sequence through AF2 takes only 5-10 sec.

During each iteration of the genetic algorithm, the sequences of the population are run through AF2 and the predicted structure along with other AF2 outputs such as confidence metrics are collected to use for scoring. In the design simulations, we run the sequences of the miniprotein through AF2 both as a monomer and with HA-PD1 as a complex and use both predictions for scoring.

### ProteinMPNN

We use a local installation of ProteinMPNN to generate new sequences for the pool once every 10 iterations. As ProteinMPNN runs in just ∼1 second per sequence, we do not perform any further runtime optimization to run it in a high-throughput manner. A PDB-formatted string from the AF2 prediction of each sequence in the population is used as backbone input and a single new sequence per PDB is predicted with ProteinMPNN using a sampling temperature of 0.1.

### EvoPro

The EvoPro pipeline is a genetic algorithm-based protein optimization framework written in Python 3.8 that interfaces with AF2 during the scoring step and ProteinMPNN during the pool refill step. Each iteration starts with a population of design sequences that are scored for fitness by running each sequence through AF2 for structure prediction. Based on AF2 outputs, the sequences are scored using a custom scoring function (described below). The sequences are then ranked from best score to worst, and the worse scoring half of the population is discarded. New sequences are generated to refill the pool to its original size either through 6-8 random point mutations (12.5% of the sequence) and crossovers of the remaining pool or using ProteinMPNN once every 10 iterations (as described above). This is used as the sequence pool for the next iteration and the process is repeated. After several iterations, as specified by the user, the process is halted before the next pool refill step and the sequences in the remaining half of the final pool, along with their AF2 prediction results, are written to disk as design outputs.

### EvoPro scoring

The score function used to evaluate miniprotein binders consists of three individual score components, summed using weights to calculate each sequence’s overall fitness score.

1. The placement confidence score evaluates the number of interface contacts made using the distances between sidechains at the interface between the target protein and designed minibinder in the AF2 complex prediction. If any two atoms in a pair of interchain sidechains are within the distance cutoff of 4Å, it is counted as a contact. Each contact is then weighted by the normalized PAE, which is calculated as follows:

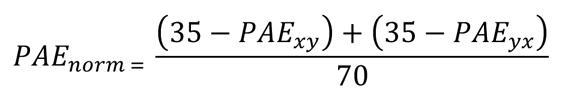 where *PAE*_*xy*_ is the PAE of residue x with respect to residue y. The total sum of weighted contacts makes up the placement score.
2. The fold confidence score is calculated by averaging the AF2 per-residue confidence score (pLDDT), which is a measure of AF2’s confidence in its backbone prediction of the residue, over the residues of the miniprotein in the AF2 prediction of the monomeric binder. We incorporate this score term to evaluate the ability of the miniprotein to fold stably in the absence of the target protein.
3. The conformational stability score is calculated as the RMSD of the carbon-alpha atoms of the miniprotein in the AF2 predictions of the target-binder complex and the monomeric binder. We incorporate this score term to minimize the energy barrier of conformational change required for the miniprotein to bind the target protein.

The overall fitness score is then the weighted sum of these 3 components:

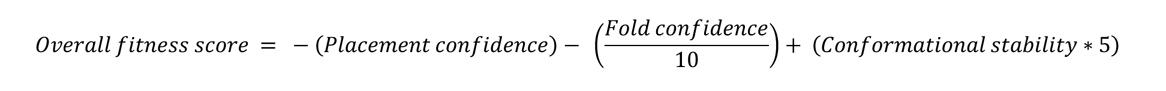

The weights were assigned based on the order of magnitude of the raw score term values, to give them relatively equal importance in the overall score. Since lower, more negative overall scores are better, score terms being maximized are negated and minimized score terms are kept as positive.

### Binder design with EvoPro

Using the pipeline described above, we ran EvoPro trajectories starting with 51 scaffolds of diverse topologies, selected from the published set of miniproteins by Cao *et al.*^1^. For each scaffold, we ran 5 independent design trajectories, each with a pool size of 50 sequences and running the genetic algorithm over 100 iterations. Since the number of outputs is half the size of the population pool, each trajectory generates 25 designs. Therefore, with 25 × 5 = 125 designed binders per starting scaffold, we generated a total of 125 × 51 = 6,375 designs.

The resulting 6,375 sequences from the EvoPro design process were orthogonally tested for interface quality using Rosetta. Using the FastRelax mover, clashes were reduced and rotamer energies were minimized. Subsequently, the InterfaceAnalyzer mover was applied to calculate an energy score for the interface. We used the calculated free energy of the interface normalized by the interface size (dG/dSASA), as a metric to sort and filter the designs to select the final sequences to be experimentally tested. Additionally, we filtered designs using an RMSD cutoff of 3Å between our design models and an OmegaFold predicted structure of the target protein linked to each AiD with a 28 amino acid glycine-serine linker for additional orthogonal verification. Finally, we applied a redundancy filter to ensure that our final set of 23 designed miniprotein sequences only contained a maximum of three designs per starting scaffold.

Upon selecting the 23 best designs from the design campaign for expression, purification, and testing, we optimized the linker length to generate the masked antagonist constructs. We predicted structures using OmegaFold of the C-terminus of the target linked to the N-terminus of each AiD with varying flexible linker lengths of 14, 16, 18, 20, 22, 24, 26, and 28 amino acids. Each linker contains the 7-residue TEV cleavage motif in the center surrounded by glycine and serine residues on either side. For each AiD, the minimum linker length that does not change the placement of the AiD in the target pocket was chosen.

### Masked antagonist mammalian expression and purification

Masked antagonists (consisting of HA-PD1, a cleavable linker, the AiD, and a C--terminal 8xHis tag) were codon optimized for mammalian systems and cloned into the mammalian expression vector pαH using the KpnI and XhoI restriction enzyme sites (Twist Bioscience) as previously described(32, 35). Masked antagonists were then expressed in Expi293F mammalian cells (Thermo Fisher Scientific, A14527) by transient transfection using the ExpiFectamine293 system (Thermo Fisher Scientific, A14525) according to a modified manufacturer’s protocol at two production scales: either for a 24-well plate or for a 125 mL flask; transfection and enhancement using Hyclone Cell Boost 1 (GE Healthcare/Cytiva, SH30584.02) as described previously(32, 35). Following expression, protein was purified using nickel affinity chromatography and size exclusion chromatography as previously described(32). Modifications specific for the 24-well plate (WHATMAN, 77015102) scale include an expression volume of 4 mL (instead of 2.5 mL), harvesting the cells by centrifugation at maximum speed for 10 min, batch binding with 150 µL nickel resin at 250 rpm for 1 h, repeating three cycles of washing with 500 µL of wash buffer followed by a spin at 250 g at 4°C for 1 min, and repeating three elution cycles of 200 µL into a 24-deep well V-bottom plate (VWR/Axygen, 89080-534), passing the previous eluant through the subsequent elution cycle.), passing the previous eluant through the subsequent elution cycle.

### PD-1:PD-L1 neutralizing activity by PD-1:PD-L1 blockade bioassay

The activity of the masked antagonists was analyzed using a PD-1/PD-L1 Blockade Bioassay (Promega, J1250) according to a modified manufacturer’s protocol as described previously(32). Briefly, PD-L1 aAPC/CHO-K1 cells were seeded into a 96-well assay plate and incubated for 16 h. For all cleaved masked antagonists, each construct (4 μM) was incubated with 75% molar equivalent TEV in assay buffer at 37°C for 16 h. The non-cleaved masked complexes (4 μM) were incubated at 4°C for 16 h.

Masked complexes ± TEV (either at 10 and 100 nM or 1:2.5 serial dilutions from 1.3 nM to 2 μM – for screening or for the full assay, respectively) were added to the plates followed by seeding PD-1 effector cells. After co-culture for 6 h, the Bio-Glo™ reagent was added. The luminescence signal was measured using a CLARIOstar *Plus* microplate reader (BMG Labtech). The full assay was performed twice either in duplicate or triplicate (technical replicates) and relative luminescence units (RLU) were plot against masked complexes concentration. The data from each experiment were analyzed using the non-linear fit in Prism Version 9.4.1 with all samples constrained to the same top and bottom value (as determined for NoMA without constraints). The data across experiments were then averaged.

### Masked antagonist surface plasmon resonance

Experiments were conducted using a Biacore 8K instrument (Cytiva) at 25°C in PBS containing 0.05% Tween 20, pH 7.4 (PBST) as previously described(32). Briefly, biotinylated PD-L1 (Sino Biological, 10084-H08H-B, reconstituted according to manufacturer’s protocol) was diluted to 500 ng/mL in PBST and immobilized onto a Series S NeutrAvidin (NA) Sensor Chip (Cytiva, 29407997) to a response (RU) of ∼100. Following immobilization, affinity measurements were made with five 1:3 serial dilutions of each recombinant masked antagonists ± TEV in PBST (incubated overnight) using a single-cycle kinetics method. All data were analyzed with the Biacore Insight Evaluation software version 3.0.12 with the general single-cycle kinetics method and a 1:1 binding kinetics fit model. In addition, data were imported to Prism version 9.4.1 to analyze data (determine the mean ± SEM across replicates) and create figures.

### AiD bacterial expression and purification

Genes coding for nine selected autoinhibitory domains (AiD4, AiD5, AiD7, AiD9, AiD10, AiD15, AiD16, AiD19, and AiD20) alone (not in complex with HA-PD1) were synthesized and cloned into pET21(+) expression vector (Twist Bioscience). An additional N-terminal sequence containing a ribosome binding site, start codon, 8xHis-tag, TEV cleavage site, and glycine spacer was added to aid purification (full sequence: GAAATAATTTTGTTTAACTTTAAGAAGGAGATATACCATGAGCCACCATCATCATCATCATCATCATTCTGAAAACTTGTATTTTCAATCGGGTGGGGGG), similar to other purification protocols^87^. These constructs were transformed into and expressed in Escherichia coli BL21(DE3)pLysS cells. Briefly, starter cultures were grown overnight at 37 °C in lysogeny broth (LB) supplemented with ampicillin. The overnight cultures were used to inoculate either 100 mL (for screening) or 1 L (for further experimental characterization) of fresh terrific broth (TB) media with an additional 8 mL 50% glycerol solution supplemented with ampicillin. Cells were grown at 37 °C until an optical density measured at a wavelength of 600 nm (OD600) of 1.0–1.2, upon which the temperature was decreased to 16°C and protein production was induced via the introduction of isopropyl β-D-1-thiogalactopyranoside to 600 μM. Cells were grown overnight at 16 °C at 200 rpm for typically 18 to 20 h. Cells were harvested by centrifugation, the cell pellets were solubilized and incubated at 25°C for an hour in lysis buffer (500 mM NaCl, 50 mM Tris HCl pH 7.5, 10 mM Imidazole, 25% B-PER solution (Thermofisher, 90084), 2.5 mg/mL lysozyme (Thermofisher, 89833)) which was mixed with protease inhibitors (phenylmethylsulfonyl fluoride, Pepstatin A, Bestatin (Thermofisher, 78433), Leupeptin (Sigma-Aldrich, L2023-100MG)), and *Serratia marcescens* nuclease A. Cell lysates were clarified by centrifugation (15,000 × g for 30 min) and filtered (5.0 μM nylon membrane) before loading onto an Ni-NTA gravity flow column for affinity chromatography. The column was washed with 20 volumes of 25 mM imidazole, PBS pH 7.4 followed by 20 volumes of 50 mM imidazole, PBS pH 7.4 before elution with 500 mM imidazole, PBS pH 7.4.

### Recombinant AiD affinity for HA-PD1 via surface plasmon resonance

SPR experiments were performed with a Biacore 8K instrument (Cytiva) using PBST. For binding assays of AiDs with HA-PD1, HA-PD1 with a C-terminal AviTag was biotinylated with BirA according to the manufacturer’s protocol (Avidity) and immobilized on a NeutrAvidin chip (Cytiva, 29407997) to a level of ∼250 response units (RUs). Binding measurements were performed using single-cycle kinetics with five 1:3 serial dilutions of each recombinant AiD (at concentrations indicated in figures) using 120 seconds of association and 1200 seconds for the final dissociation time at 25°C and a flow rate of 30 μL/min. The data were analyzed in the same manner as the masked antagonists.

### AiD concentration determination

Because the AiDs contained few tyrosine or tryptophan residues, the protein concentrations were determined by image analysis of SDS-PAGE with 4-20% Mini-PROTEAN® TGX Stain-Free™ Protein Gels (Bio-Rad, 4568096) followed by Coomassie Brilliant Blue staining using the following concentration standard: Precision Plus Protein™ Unstained Protein Standards, Strep-tagged recombinant (BioRad, 1610363). The gel image was acquired with ChemiDoc Imaging Systems (BioRad) and analyzed with ImageLab software (BioRad).

### Circular dichroism (CD)

All CD experiments were performed with a Jasco J-815 CD spectrometer. Data were collected in PBS pH 7.4 buffer at AiD protein concentrations ranging from 10 to 40 µM. CD spectra were collected from 250 to 190 nm at 20°C in 1 mm path-length quartz cuvettes using the following collection parameters: 8 second data integration time (DIT), 2 nm band width, 1 nm data pitch, 50 nm/min scanning speed, and 3 accumulations. Thermal denaturation experiments were conducted similarly except that: 1) signal collected at 222 nm, 2) temperature ranging from 5 to 95°C, 3) 2°C/min ramp rate, 4) 16 second DIT, and 5) the temperature ramp was paused while collecting data.

#### Abbreviations

2H3E: two helices with three ý-strands
2H4E: two helices with four ý-strands
3H: three-helix bundle
4H: four-helix bundle
AF2: AlphaFold2
AiD: autoinhibitory domain
CD: circular dichroism
CDR: complementarity-determining regions dG/dSASA dG_separated/dSASA × 100
DL: deep learning HA-PD1 high affinity PD-1
ipTM: interface predicted template modeling score K_D_ equilibrium dissociation constant
k_off_: rate constant of dissociation
k_on_: rate constant of association
MA: masked antagonist
NoMA: antagonist without a mask MSA multiple sequence alignment
PAE: predicted aligned error
PD-1: programmed cell death receptor 1 PD-L1 programmed death-ligand 1
pLDDT: predicted local distance difference test RMSD root mean square deviation
SPR: surface plasmon resonance
TCR: T-cell receptor
TEV: Tobacco Etch Virus

## Contact and competing interest information

Odessa J. Goudy

Amrita Nallathambi

Tomoaki Kinjo

Nicholas Randolph

Brian Kuhlman

None of the authors have any competing interests.

## Data sharing plans

Python scripts for EvoPro can be found at https://github.com/Kuhlman-Lab/evopro.

