## Supplementary materials for "*In silico* evolution of protein binders with deep learning models for structure prediction and sequence design"

### Supplementary Information

Supplementary Tables: Table S1-S4

**Supplementary Table 1.** Starting miniprotein sequences

**Supplementary Table 2.** EvoPro-designed binder sequences

**Supplementary Table 3.** Masked antagonist protein sequences

**Supplementary Table 4.** Recombinant AiD kinetic parameters

Supplementary Figures: Figure S1-S12

**Supplementary Figure 1.** HA-PD1 binding groove pocket

**Supplementary Figure 2.** Masked antagonist expression screen

**Supplementary Figure 3.** Masked antagonist cellular activity screen

**Supplementary Figure 4.** Masked antagonist cleavage during cellular assay

**Supplementary Figure 5.** Masked antagonist expression and purification

**Supplementary Figure 6.** Masked antagonist cellular assay

**Supplementary Figure 7.** SPR sensorgrams of masked antagonists

**Supplementary Figure 8.** Recombinant AiD expression

**Supplementary Figure 9.** CD spectra of recombinant AiDs

**Supplementary Figure 10.** SPR sensorgrams of recombinant AiDs

**Supplementary Figure 11.** Mutation of key AiD interface residues

**Supplementary Figure 12.** SPR sensorgrams of mutant recombinant AiDs

| Scaffold | Sequence | Topology |
|---|---|---|
| 1 | SIDEELERLLEEAGVDPELIDDLYAVIYQLYIRGLSDKDVLRFLLENLERDGTPLRRVVEELLKR | 4H |
| 2 | RDEIKERIFKAVVRAIVTGNPEQLKEAKKLLEKLKKLGRLDQDAKKFEKAIRQVEKRLR | 3H |
| 3 | NAEEITEKATLVGIEAWLLAKDEEQKKKVRTLNRQVKKLLQQNDLDQAKRVLDQLKSVLEDLK | 3H |
| 4 | DRRKEMDKVYRTAFKRITSTPDKEKRKEVVKEATEQLRRIAKDEEEKKKAAYMILFLKTL | 3H |
| 5 | GGGDRRKEMDKVYRTAFKRITSTPDKEKRKEVVKEATEQLRRIAKDEEEKKKAAYMILFLKTL | 3H |
| 6 | SVIEKLRKLEKQARKQGDEVLVMLARMVLEYLEKGWVSEEDADESADRIEEVLKK | 3H |
| 7 | NHIACEIHNPEAAKEIAKVANVRRVYFIKQPGNRYFVLLKNADPEGVKKVRSKYNVRCVIRE | 2H4E |
| 8 | SVIEKLRKLEKQARKQGDEVLVMLARMVLEYLEKGWVSEEDADESADRIEEVLKK | 3H |
| 9 | DEELMELVYEAVEKNDPELLFEAWMELASLLDETGDPKIEEALGLLQQVDGGNPDAGRRIKELFK | 4H |
| 10 | DDERLATLAFRALIKRAGVKNLDVKVTNGKVRVTITGRDQASFKALQLVFALARRLGLQVQIDTR | 2H3E |
| 11 | HCTIEVVGVDPEKVEAIAAAYGAEVCEKDGKFEIHLDDPHSAESAAVAISVLTNRPVRLQC | 2H4E |
| 12 | DTQTLIKIVTDLAREVIKRVQDEKQQRRAKTLAKQVQKSRDPELAKTVSKELKDILKST | 3H |
| 13 | GPELEKKARKLLEEAQKKQDWKLADKAVKVAEKVLKEVGNPELEKLLRTAQTVRKQIKG | 3H |
| 14 | SLKDKAKSLYESATKSVKNASSEEEKKRILDTAVSELNKIPDPTAQKLSEKLERVWRSLS | 3H |
| 15 | SQKDKAKSLYESATKSVKHASSEEEKKRILDTAVSELNKISDPTAQKLSEKLERVWRSLS | 3H |
| 16 | SLKDKAKSLYESATKSVKHASSEEEKKRILDTAVSELNKIPDPLAQKLSEKLERVWRSLS | 3H |
| 17 | ETKEVTFDNPEEATKFAQKLGNEEAVRIAVKLAKELQRPVTVRVNNKVEVTVNGDRVTIREHS | 2H4E |
| 18 | GTLKITVRNLDPNLVKTAAKIVNVKVEVHGQTVTLKDASEELAKKFQEVAKRLGLDLEVSVEF | 2H4E |
| 19 | EVTVEFRNASPEAIKLLVKLVNAESYEVHGNTVHLEGVDPRRVKEAAKIVSKTFNVEVHVRG | 2H4E |
| 20 | TVQLHVKNADPKQVEEAAKIVGVRVEIKPGNQIDIDVSDPSLVDIVAKLANVSEWEVRG | 2H4E |
| 21 | GTVTFHGVDPNVVKEAVSKAGAEVRVHGETVEVHVSDDNLAKKVLELVRKLAGAEVEIHS | 2H4E |
| 22 | DKEEEELKRVVERLERLNPEIREVLRTVLELLKRDPEEARRFLKEIVEKTGDPLLKELLEVLEET | 4H |
| 23 | SEEEVRRILREAVEKNDPRLLRKAFELLERLLGDPEEALKVLIRILEELGVFSPEELEELKRLLR | 4H |
| 24 | DEEKIEEAIKKAEEEGNPEVALVLRLLLELLKRFPPEEVEEIAEKIAKELGIPPEVIKRAKELFR | 4H |
| 25 | DEKEELLRELQEKIPDPRVREVLELAIKLLEDGVPPEEIKKLIEKLVKELGLPPEVLELVEKIVK | 4H |
| 26 | TKEKLREIVRLLREGDPRARELLERLLKELRERGNEEEVRIVEKILELLKRDPEKATRLLEKLAR | 4H |
| 27 | DKEEAERLLERVKRIEDPKERRELLERAEELARRANDPELLREIEEELRKLN | 3H |
| 28 | DLEEAVRKLEELLRRGDPEKVEKLRKELEEILKKVTDPELRREAERVLREAEEFLRR | 3H |
| 29 | DVEEYLEEIKRRLEELRKKGDEEELERLTREAEEYLERKGNPELVEELRRFLEELG | 3H |

| | | |
|---|---|---|
| 30 | DEEIERLERELRRLKEEGNEEELKKLLEEIEELVRRTGDPRVKELLKLARKALEEL | 3H |
| 31 | DEEEERVRRFAEEVERLTKDPELQEEVRELLERYEREGDPRYVELIEKIYEEVKK | 3H |
| 32 | DVTITVKNPEVAKNASKIAGVQSYEIEEHGDRITVHLHGVDPRLARKAEKYVRSVVPDAEIHVS | 2H4E |
| 33 | DTKWTLTFDDPEAAKVVASLVGAEVRTHNNKKVTVHLDSSSLVELAKEVSKKLGVPFEVREETS | 2H4E |
| 34 | TVTIHVRNPTEEVKKAAEYAQSYGVEVHFHGPDIELRNVDPEIAKTVYELIQKAGAEEVHVTIY | 2H4E |
| 35 | NVTVHIHNADPNLVKSAASSAGEVEVEYKDGTIKVHVKDPNLAKKVSDIIRKLVGAEVHVQQN | 2H4E |
| 36 | DFTFNVSLDPKLVKKVASYAGAEVHVEKHGDTVQVHLKGADPTLVKEVEKLLRKLAGSVEVRSS | 2H4E |
| 37 | QQKYSVTVDDPHAAKLAQEVAQRLGAEYHVKDDKTVTVTVPSKELAKKLQKLVGFPVEIRTVQD | 2H4E |
| 38 | PVTITVTIDDVKKAERAAKIVGAEVHVKREGDEWKVTVHVENEELAKLVSKLAGVKDFRIHSG | 2H4E |
| 39 | PKYTVQVHADPTEVKKAVSKAGAKIETEPGNQVKVHVDDPSIAKTLETLLKQLGVEFEIRSRE | 2H4E |
| 40 | GKVEVHVDDVDRVKEAVKKANATVKVHGKKVEVEIESEEVSKKVSKVLKEAGVKSFSVQF | 2H4E |
| 41 | KTVDINVTGADPTIASELAKEVNARVETHGETITLKLHDVDPRLAKKVVDSLRNWNAEVHVQF | 2H4E |
| 42 | DREEAKERIKELLELVKRVSEEERRELLREARRLAERVNDPEARRLVEELERLIKEL | 3H |
| 43 | DLEERAKEVLERIEEAIERNDPETVEKLLRELKELAKRLNDEELKRFIEEVEKWVK | 3H |
| 44 | SEEELERLEREIRRLLKEGDLEEAKRLIEELKRLAEELNDEEAKEIVKELEEILRKL | 3H |
| 45 | SEEELERLLRELEEALRRGDEELFEKLREEVEKLARELNDEEVLKRIEELERLLH | 3H |
| 46 | DEERVRELLEEAKRLAKEGDPEEARKLLEEAKRIAEESNDEELERLIREVEEEVKEL | 3H |
| 47 | DVEERVKKLLEKAKHIPDPEERRRVLEEARRLAEELNDPELEKLVEEVERRL | 3H |
| 48 | DEVEREAEEVERLAREKNNPELLREVERLLKELREKGDPELERELRELLEKVKRLL | 3H |
| 49 | SEEVEEAEELAEEAERLVRELGDEELLRELEELLKRLREEGDPRLAEELRRVAKRIL | 3H |
| 50 | SLVERVRELLERALKRNDPELRREAEEFAEEVLRKLKDERLRERVERLLREFKKRI | 3H |
| 51 | DVEELARRLLERLKKEGNEEEFEKLRRLLEKLEREGDPRLREELRRLIEEIN | 3H |

**Supplementary Table 1. Starting scaffold miniprotein sequences for EvoPro runs.**
These protein sequences were used as input to EvoPro. Each sequence was assigned a random ID number. All sequences were obtained from a database of *de novo* proteins characterized for hyperstability (Cao, L., Coventry, B., Goreshnik, I. *et al.* Design of protein-binding proteins from the target structure alone. *Nature* **605**, 551–560 (2022). https://doi.org/10.1038/s41586-022-04654-9).

| | EvoPro design | | Starting scaffold miniprotein | |
|---|---|---|---|---|
| AiD | Sequence (AiD only) | dG/ dSASA | Sequence | Sequence identity (%) |
| 0 | TVEFRVRLPPEVAQEICDAAGVEARLRVEGDTVLVTLRGASEEQVAAVEAALRAFAGQVEREEL | -4.258 | DFTFNVSLDPKLVKKVASYAGAEVHVEKHGDTVQVHLKGADPTLVKEVEKLLRKLAGSVEVRSS | 37.5 |
| 1 | AEELRKEALELMQKAVEEKNAELAAEAVKIAQKAYDLTGDTRSQLLLDGAQILLQDLTA | -4.185 | GPELEKKARKLLEEAQKKQDWKLADKAVKVAEKVLKEVGNPELEKLLRTAQTVRKQIKG | 30.5 |
| 2 | SDEEKLMEILRILQEALTVGPEERKKLCEEARELAKKVSDPMRRMLVLYAIKLVLNL | -4.168 | DREEAKERIKELLELVKRVSEEERRELLREARRLAERVNDPEARRLVEELERLIKEL | 38.6 |
| 3 | DAELEAVRAVAQAVIAESDDPEIRAFARRLLAQYKATGNGIYADALRLLLRDVQA | -4.158 | DEEEERVRRFAEEVERLTKDPELQEEVRELLERYEREGDPRYVELIEKIYEEVKK | 30.9 |
| 4 | SDEEKLMEILRILQEALTVGPEERALLCEEARELAEQISDPMRRQLVLYAIKLVLNL | -4.080 | DREEAKERIKELLELVKRVSEEERRELLREARRLAERVNDPEARRLVEELERLIKEL | 38.6 |
| 5 | SDEEKIMEIEKILQEALTVGPEERALLCEEARELAEQISDPMRRMLVLYAIKLVLNF | -4.041 | DREEAKERIKELLELVKRVSEEERRELLREARRLAERVNDPEARRLVEELERLIKEL | 36.8 |
| 6 | RTYYLVTFPTHEVAVEAAAKVGGTVIRLPDGSGLLILDREEDVARALAVGAAAGVPATVERIEV | -4.030 | DTKWTLTFDDPEAAKVVASLVGAEVRTHNNKKVTVHLDSSSLVELAKEVSKKLGVPFEVREETS | 28.1 |
| 7 | NPELRALVREAAETRSLNLLLDALIKVYKEYQKTGDPRLERILELLAAVIEGDEEAREELLALLA | -3.991 | DEELMELVYEAVEKNDPELLFEAWMELASLLDETGDPKIEEALGLLQQVDGGNPDAGRRIKELFK | 33.8 |
| 8 | SLFQEIMDIYNEALEKAEKAKSTEEKIAILDEAIEKLRKIDDYAARVVADYLEEEREKLL | -3.972 | SLKDKAKSLYESATKSVKHASSEEEKKRILDTAVSELNKIPDPLAQKLSEKLERVWRSLS | 35.0 |
| 9 | SKEEVLELLQDALELAKEGKIEYAFANVKEARKLAEELGDGILLDAVYLVELEIEEL | -3.966 | DEERVRELLEEAKRLAKEGDPEEARKLLEEAKRIAEESNDEELERLIREVEEEVKEL | 439. |
| 10 | TREEIETLLDLALREARRGNFEEANKLLEEARKIAEELGDGLMLYTIELVRKEVEAL | -3.957 | DEERVRELLEEAKRLAKEGDPEEARKLLEEAKRIAEESNDEELERLIREVEEEVKEL | 45.6 |
| 11 | SMEEQIEELLENAAKSNNETSRAVYIYQAYILAKKISPEMVEYVYEAADEIL | -3.908 | DVEERVKKLLEKAKHIPDPEERRRVLEEARRLAEELNDPELEKLVEEVERRL | 25.0 |
| 12 | WRVVATGARPAAVEEVAAAAGVPVWREGDRYVLEVPDLELARLMARAIELLGGGTAVVEP | -3.895 | GTVTFHGVDPNVVKEAVSKAGAEVRVHGETVEVHVSDDNLAKKVLELVRKLAGAEVEIHS | 25.0 |
| 13 | MAVVRARGADPRVVREIAAAVGPAVVEEEDGVDTVRMEDPVKGALIADAIRIATGAEVDYEVL | -3.874 | NVTVHIHNADPNLVKSAASSAGEVEVEYKDGTIKVHVKDPNLAKKVSDIIRKLVGAEVHVQQN | 33.3 |
| 14 | SLEEVLEILRRAIEEDDPELVVEAYEQLVRMTGSRTLALEILRQLLEELGCTSPRLWATVDALRA | -3.873 | SEEEVRRILREAVEKNDPRLLRKAFELLERLLGDPEEALKVLIRILEELGVFSPEELEELKRLLR | 43.1 |
| 15 | TTTTKTFTDLDEALAYADKIGIYSAVLYAYARSLATGKPVTVNAGGGYTVTVRDGVITITKTT | -3.853 | ETKEVTFDNPEEATKFAQKLGNEEAVRIAVKLAKELQRPVTVRVNNKVEVTVNGDRVTIREHS | 31.7 |
| 16 | DLLERLAALLTARGVPADERQGVLELVAYLLRWGQSWEWALRFIVEDTPDPNHPVRLAAEELLAE | -3.838 | SIDEELERLLEEAGVDPELIDDLYAVIYQLYIRGLSDKDVLRFLLENLERDGTPLRRVVEELLKR | 29.2 |
| 17 | TVYRVTVDADTPELREAARALGGEVERLPGGVLVVYVTKALAADGLRAMARSLGATARIEALP | -3.831 | PKYTVQVHADPTEVKKAVSKAGAKIETEPGNQVKVHVDDPSIAKTLETLLKQLGVEFEIRSRE | 28.6 |
| 18 | SLVDLVRIVLEIALESKAPELQAQAYALASHIIARSRDPYVRARIEALLLDFEARL | -3.810 | SLVERVRELLERALKRNDPELRREAEEFAEEVLRKLKDERLRERVERLLREFKKRI | 39.3 |
| 19 | SARVERFRAEAAELPDPAARELMAEAEELYLAGVNPSLILLAVRIAVRERGLPPEVVALVEELVA | -3.804 | DEKEELLRELQEKIPDPRVREVLELAIKLLEDGVPPEEIKKLIEKLVKELGLPPEVLELVEKIVK | 38.5 |
| 20 | SKEELEVYMRYLEWALRRGDEVLFQKLFEETVKLASELNDNVILREVALLDYLWH | -3.798 | SEEELERLLRELEEALRRGDEELFEKLREEVEKLARELNDEEVLKRIEELERLLH | 58.2 |
| 21 | SAREEVIALLEQARAEPDKARALEILDRAEALAREIGDGLLVEAVRLLRREI | -3.775 | DVEERVKKLLEKAKHIPDPEERRRVLEEARRLAEELNDPELEKLVEEVERRL | 32.7 |
| 22 | LQEVIEHLKLLITRAEEYKTIEILEEAEELYRYLNRLGKLGEEGEEIRERIEEVENEIS | -3.760 | RDEIKERIFKAVVRAIVTGNPEQLKEAKKLLEKLKKLGRLDQDAKKFEKAIRQVEKRLR | 27.1 |

**Supplementary Table 2. EvoPro-designed binders selected for further experimental characterization.** Autoinhibitory domains (AiDs) designed using the EvoPro pipeline (left) are compared to the original starting sequence (right). Each AiD sequence was assigned an identification number based on the dG/dSASA rank.

| MA | Sequence |
|---|---|
| 4 | WNPPTFSPALLVVTEGDNATFTCSFSNTSESFHVVWHRESPSGQTDTLAAFPEDRSQPGQDSRFRVTQLPNGRDFHMSVVRARRNDSGTYVCGVISLAPKIQIKESLRAELRVTERRAEGGSGSGGENLYFQGGSG<br>SGG**SDEEKLMEILRILQEALTVGPEERALLCEEARELAEQISDPMRRQLVLYAIKLVLNL** |
| 5 | WNPPTFSPALLVVTEGDNATFTCSFSNTSESFHVVWHRESPSGQTDTLAAFPEDRSQPGQDSRFRVTQLPNGRDFHMSVVRARRNDSGTYVCGVISLAPKIQIKESLRAELRVTERRAEGGSGSGGENLYFQGGSG<br>SGG**SDEEKIMEIEKILQEALTVGPEERALLCEEARELAEQISDPMRRMLVLYAIKLVLNF** |
| 7 | WNPPTFSPALLVVTEGDNATFTCSFSNTSESFHVVWHRESPSGQTDTLAAFPEDRSQPGQDSRFRVTQLPNGRDFHMSVVRARRNDSGTYVCGVISLAPKIQIKESLRAELRVTERRAEGGSGGSGGENLYFQGGS<br>GGSGG**NPELRALVREAAETRSLNLLLDALIKVYKEYQKTGDPRLERILELLAAVIEGDEEAREELLALLA** |
| 9 | WNPPTFSPALLVVTEGDNATFTCSFSNTSESFHVVWHRESPSGQTDTLAAFPEDRSQPGQDSRFRVTQLPNGRDFHMSVVRARRNDSGTYVCGVISLAPKIQIKESLRAELRVTERRAEGGSGSGGENLYFQGGSG<br>SGG**SKEEVLELLQDALELAKEGKIEYAFANVKEARKLAEELGDGILLDAVYLVELEIEEL** |
| 10 | WNPPTFSPALLVVTEGDNATFTCSFSNTSESFHVVWHRESPSGQTDTLAAFPEDRSQPGQDSRFRVTQLPNGRDFHMSVVRARRNDSGTYVCGVISLAPKIQIKESLRAELRVTERRAEGSGGGENLYFQGGGGSG<br>G**TREEIETLLDLALREARRGNFEEANKLLEEARKIAEELGDGLMLYTIELVRKEVEAL** |
| 15 | WNPPTFSPALLVVTEGDNATFTCSFSNTSESFHVVWHRESPSGQTDTLAAFPEDRSQPGQDSRFRVTQLPNGRDFHMSVVRARRNDSGTYVCGVISLAPKIQIKESLRAELRVTERRAEGSGGGENLYFQGGGGSG<br>G**TTTTKTFTDLDEALAYADKIGIYSAVLYAYARSLATGKPVTVNAGGGYTVTVRDGVITITKTT** |
| 16 | WNPPTFSPALLVVTEGDNATFTCSFSNTSESFHVVWHRESPSGQTDTLAAFPEDRSQPGQDSRFRVTQLPNGRDFHMSVVRARRNDSGTYVCGVISLAPKIQIKESLRAELRVTERRAEGSGGGENLYFQGGGGSG<br>G**DLLERLAALLTARGVPADERQGVLELVAYLLRWGQSWEWALRFIVEDTPDPNHPVRLAAEELLAE** |
| 17 | WNPPTFSPALLVVTEGDNATFTCSFSNTSESFHVVWHRESPSGQTDTLAAFPEDRSQPGQDSRFRVTQLPNGRDFHMSVVRARRNDSGTYVCGVISLAPKIQIKESLRAELRVTERRAEGGSGSGGENLYFQGGSG<br>SGG**TVYRVTVDADTPELREAARALGGEVERLPGGVLVVYVTKALAADGLRAMARSLGATARIEALP** |
| 19 | WNPPTFSPALLVVTEGDNATFTCSFSNTSESFHVVWHRESPSGQTDTLAAFPEDRSQPGQDSRFRVTQLPNGRDFHMSVVRARRNDSGTYVCGVISLAPKIQIKESLRAELRVTERRAEGGSGSGGENLYFQGGSG<br>SGG**SARVERFRAEAAELPDPAARELMAEAEELYLAGVNPSLILLAVRIAVRERGLPPEVVALVEELVA** |
| 20 | WNPPTFSPALLVVTEGDNATFTCSFSNTSESFHVVWHRESPSGQTDTLAAFPEDRSQPGQDSRFRVTQLPNGRDFHMSVVRARRNDSGTYVCGVISLAPKIQIKESLRAELRVTERRAEGGSGSGGENLYFQGGSG<br>SGG**SKEELEVYMRYLEWALRRGDEVLFQKLFEETVKLASELNDNVILREVALLDYLWH** |
| 0 | WNPPTFSPALLVVTEGDNATFTCSFSNTSESFHVVWHRESPSGQTDTLAAFPEDRSQPGQDSRFRVTQLPNGRDFHMSVVRARRNDSGTYVCGVISLAPKIQIKESLRAELRVTERRAEGGGSGGSGGENLYFQGG<br>SGGSGGG**TVEFRVRLPPEVAQEICDAAGVEARLRVEGDTVLVTLRGASEEQVAAVEAALRAFAGQVEREEL** |
| 1 | WNPPTFSPALLVVTEGDNATFTCSFSNTSESFHVVWHRESPSGQTDTLAAFPEDRSQPGQDSRFRVTQLPNGRDFHMSVVRARRNDSGTYVCGVISLAPKIQIKESLRAELRVTERRAEGSGGGENLYFQGGGGSG<br>G**AEELRKEALELMQKAVEEKNAELAAEAVKIAQKAYDLTGDTRSQLLLDGAQILLQDLTA** |
| 2 | WNPPTFSPALLVVTEGDNATFTCSFSNTSESFHVVWHRESPSGQTDTLAAFPEDRSQPGQDSRFRVTQLPNGRDFHMSVVRARRNDSGTYVCGVISLAPKIQIKESLRAELRVTERRAEGGSGSGGENLYFQGGSG<br>SGG**SDEEKLMEILRILQEALTVGPEERKKLCEEARELAKKVSDPMRRMLVLYAIKLVLNL** |
| 3 | WNPPTFSPALLVVTEGDNATFTCSFSNTSESFHVVWHRESPSGQTDTLAAFPEDRSQPGQDSRFRVTQLPNGRDFHMSVVRARRNDSGTYVCGVISLAPKIQIKESLRAELRVTERRAEGGSGGSGGENLYFQGGS<br>GGSGG**DAELEAVRAVAQAVIAESDDPEIRAFARRLLAQYKATGNGIYADALRLLLRDVQA** |
| 6 | WNPPTFSPALLVVTEGDNATFTCSFSNTSESFHVVWHRESPSGQTDTLAAFPEDRSQPGQDSRFRVTQLPNGRDFHMSVVRARRNDSGTYVCGVISLAPKIQIKESLRAELRVTERRAEGSGGGENLYFQGGGGSG<br>G**RTYYLVTFPTHEVAVEAAAKVGGTVIRLPDGSGLLILDREEDVARALAVGAAAGVPATVERIEV** |
| 8 | WNPPTFSPALLVVTEGDNATFTCSFSNTSESFHVVWHRESPSGQTDTLAAFPEDRSQPGQDSRFRVTQLPNGRDFHMSVVRARRNDSGTYVCGVISLAPKIQIKESLRAELRVTERRAEGSGGGENLYFQGGGGSG<br>G**SLFQEIMDIYNEALEKAEKAKSTEEKIAILDEAIEKLRKIDDYAARVVADYLEEEREKLL** |
| 11 | WNPPTFSPALLVVTEGDNATFTCSFSNTSESFHVVWHRESPSGQTDTLAAFPEDRSQPGQDSRFRVTQLPNGRDFHMSVVRARRNDSGTYVCGVISLAPKIQIKESLRAELRVTERRAEGSGGGENLYFQGGGGSG<br>G**SMEEQIEELLENAAKSNNETSRAVYIYQAYILAKKISPEMVEYVYEAADEIL** |
| 12 | WNPPTFSPALLVVTEGDNATFTCSFSNTSESFHVVWHRESPSGQTDTLAAFPEDRSQPGQDSRFRVTQLPNGRDFHMSVVRARRNDSGTYVCGVISLAPKIQIKESLRAELRVTERRAEGGSGSGGSGGGENLYFQ<br>GGGGSGGSGSG**WRVVATGARPAAVEEVAAAAGVPVWREGDRYVLEVPDLELARLMARAIELLGGGTAVVEP** |
| 13 | WNPPTFSPALLVVTEGDNATFTCSFSNTSESFHVVWHRESPSGQTDTLAAFPEDRSQPGQDSRFRVTQLPNGRDFHMSVVRARRNDSGTYVCGVISLAPKIQIKESLRAELRVTERRAEGSGGGENLYFQGGGGSG<br>G**MAVVRARGADPRVVREIAAAVGPAVVEEEDGVDTVRMEDPVKGALIADAIRIATGAEVDYEVL** |
| 14 | WNPPTFSPALLVVTEGDNATFTCSFSNTSESFHVVWHRESPSGQTDTLAAFPEDRSQPGQDSRFRVTQLPNGRDFHMSVVRARRNDSGTYVCGVISLAPKIQIKESLRAELRVTERRAEGSGGGENLYFQGGGGSG<br>G**SLEEVLEILRRAIEEDDPELVVEAYEQLVRMTGSRTLALEILRQLLEELGCTSPRLWATVDALRA** |
| 18 | WNPPTFSPALLVVTEGDNATFTCSFSNTSESFHVVWHRESPSGQTDTLAAFPEDRSQPGQDSRFRVTQLPNGRDFHMSVVRARRNDSGTYVCGVISLAPKIQIKESLRAELRVTERRAEGGSGSGGENLYFQGGSG<br>SGG**SLVDLVRIVLEIALESKAPELQAQAYALASHIIARSRDPYVRARIEALLLDFEARL** |
| 21 | WNPPTFSPALLVVTEGDNATFTCSFSNTSESFHVVWHRESPSGQTDTLAAFPEDRSQPGQDSRFRVTQLPNGRDFHMSVVRARRNDSGTYVCGVISLAPKIQIKESLRAELRVTERRAEGGSGSGGENLYFQGGSG<br>SGG**SAREEVIALLEQARAEPDKARALEILDRAEALAREIGDGLLVEAVRLLRREI** |
| 22 | WNPPTFSPALLVVTEGDNATFTCSFSNTSESFHVVWHRESPSGQTDTLAAFPEDRSQPGQDSRFRVTQLPNGRDFHMSVVRARRNDSGTYVCGVISLAPKIQIKESLRAELRVTERRAEGGSGGSGGENLYFQGGS<br>GGSGG**LQEVIEHLKLLITRAEEYKTIEILEEAEELYRYLNRLGKLGEEGEEIRERIEEVENEIS** |

**Supplementary Table 3. Masked antagonist protein sequences.**

Amino Acid sequences of the 23 masked antagonists selected for experimental characterization. Bold denotes the AiD region within the masked agonist. Sequences are arranged by numerical order and by constructs that showed protease-dependent binding in the first cellular assay screen (top) (Fig. S3) and by constructs that either did not express or did not show protease-dependent activity (bottom).

| AiD | $k_{on}$ ($M^{-1}$ $s^{-1}$) | | $k_{off}$ ($s^{-1}$) | | $K_D$ (M) | |
|---|---|---|---|---|---|---|
| | **mean** | **SD** | **mean** | **SD** | **mean** | **SD** |
| 4 | 4.01E+05 | 2.02E+05 | 1.40E-02 | 4.31E-03 | 3.82E-08 | 1.01E-08 |
| 5 | 8.96E+06 | 2.30E+06 | 7.48E-03 | 8.13E-04 | 8.76E-10 | 2.23E-10 |
| 5 E10A | 3.12E+06 | 3.38E+06 | 4.61E-02 | 4.53E-02 | 2.59E-08 | 1.73E-08 |
| 5 Q14A | 1.07E+07 | 4.67E+06 | 4.72E-02 | 2.49E-02 | 4.40E-09 | 2.26E-09 |
| 5 L46A | 2.38E+06 | 1.12E+06 | 4.08E-02 | 2.97E-02 | 1.57E-08 | 5.79E-09 |
| 5 Y49A | 4.13E+06 | 2.93E+05 | 6.28E-02 | 3.81E-02 | 1.52E-08 | 9.42E-09 |
| 7 | 8.42E+05 | 1.05E+05 | 3.85E-01 | 8.48E-02 | 4.55E-07 | 5.68E-08 |
| 15 | 7.51E+05 | 2.04E+05 | 1.01E-01 | 3.69E-02 | 1.35E-07 | 3.02E-08 |
| 15 L27A | 9.73E+04 | 3.80E+04 | 3.91E-01 | 2.48E-01 | 5.09E-06 | 5.02E-06 |
| 15 Y28A | 1.65E+05 | 5.20E+04 | 3.18E-01 | 9.44E-02 | 2.08E-06 | 8.22E-07 |
| 19 | 2.11E+06 | 7.70E+05 | 1.22E-01 | 2.78E-02 | 6.06E-08 | 1.22E-08 |
| 19 L29A | 7.62E+05 | 2.02E+05 | 6.60E-01 | 1.88E-01 | 8.64E-07 | 5.29E-08 |
| 19 L38A | 4.90E+04 | 4.97E+03 | 3.97E-01 | 5.15E-02 | 8.20E-06 | 1.56E-06 |

**Supplementary Table 4. Recombinant AiD binding parameters.**
Table of mean kinetic parameters for the various recombinant AiDs and mutants binding to HA-PD1 as measured with SPR. Error represents SD. The mean and SD values are calculated from at least three experiments for each AiD.

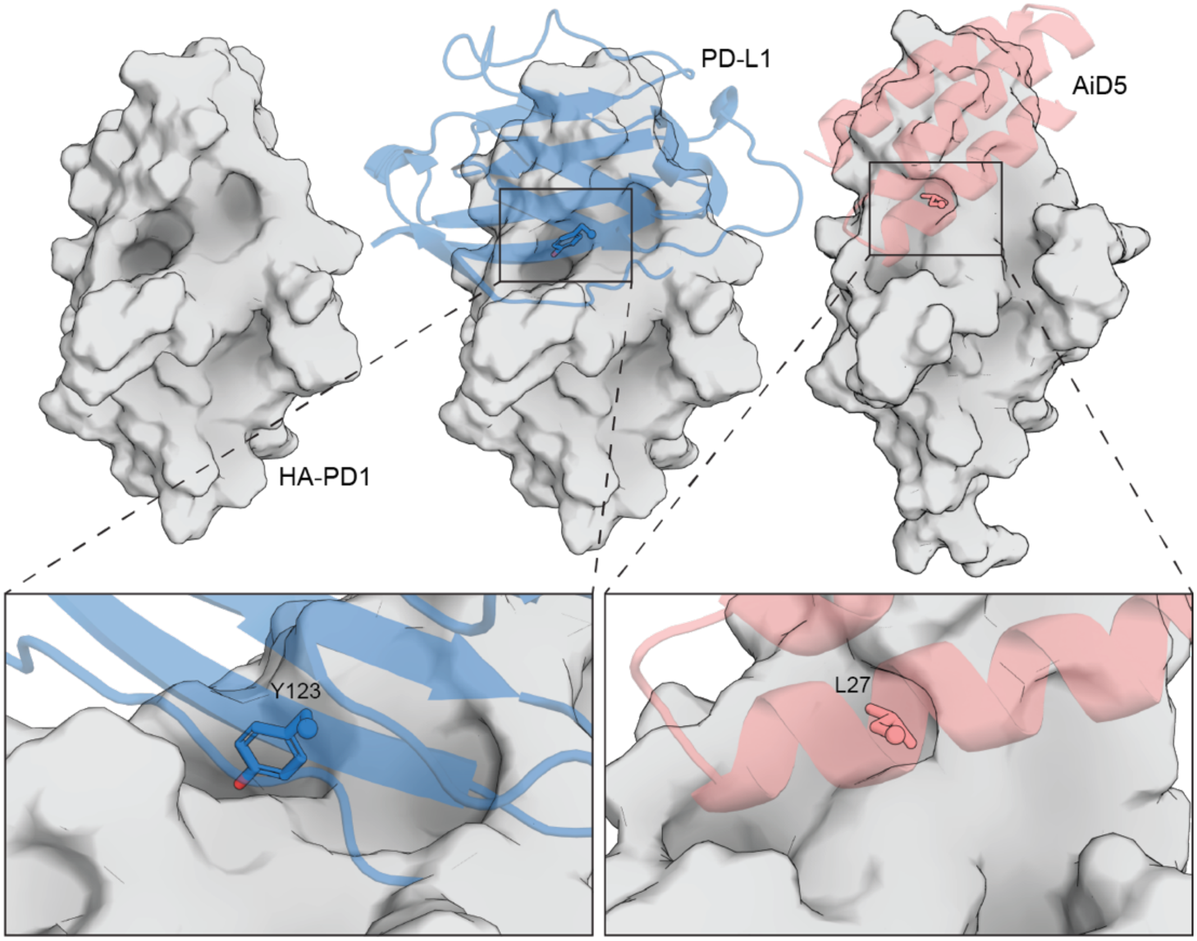


**Supplemental Figure 1. HA-PD1 binding groove has a deep pocket.**
Surface representation of HA-PD1 (grey) from the crystal structure (PDB ID: 5IUS) of the complex with native binding partner PD-L1 (blue) reveals a deep hydrophobic pocket at the binding groove that is satisfied by TYR123 (left: PD-L1 hidden, middle: PD-L1 shown). Some successful EvoPro designs position other hydrophobic residues such as isoleucine and leucine to fill the pocket. AF2 model of the target-binder complex (right) shows AiD5 (salmon) is predicted to place LEU27 in the pocket when binding. Figure created in PyMOL.

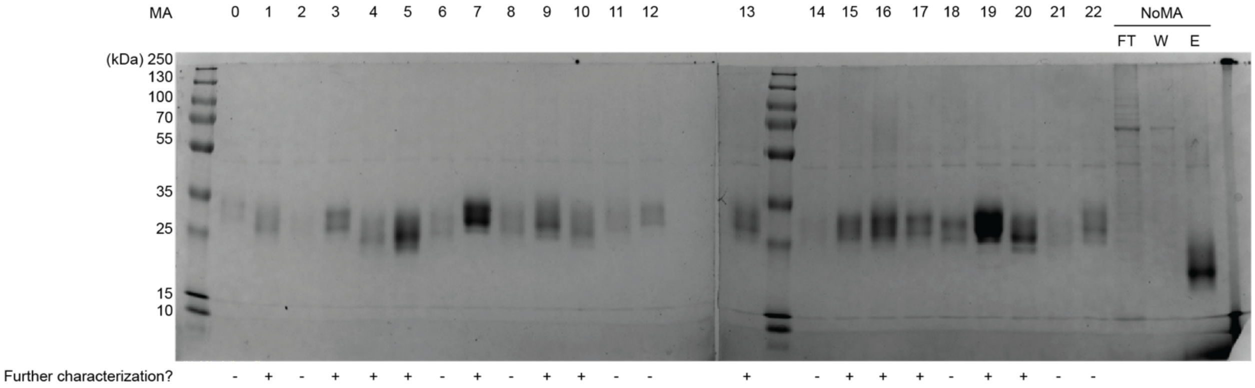


**Supplemental Figure 2. Masked antagonist expression screen.**
SDS-PAGE (16% acrylamide) of the 23 masked antagonists after expression in a 24-well plate format and the Nickle-His purification process. As a positive control, NoMA was purified in the same manner, which eluted in the elution (E) fraction. Smearing is due to glycosylation from the Expi293F mammalian expression system. We carried forward a subset of 13 well-expressing masked antagonists (denoted by a "+" below the gel) for further characterization based on a combination of the expression level and scaffold diversity of the AiD.

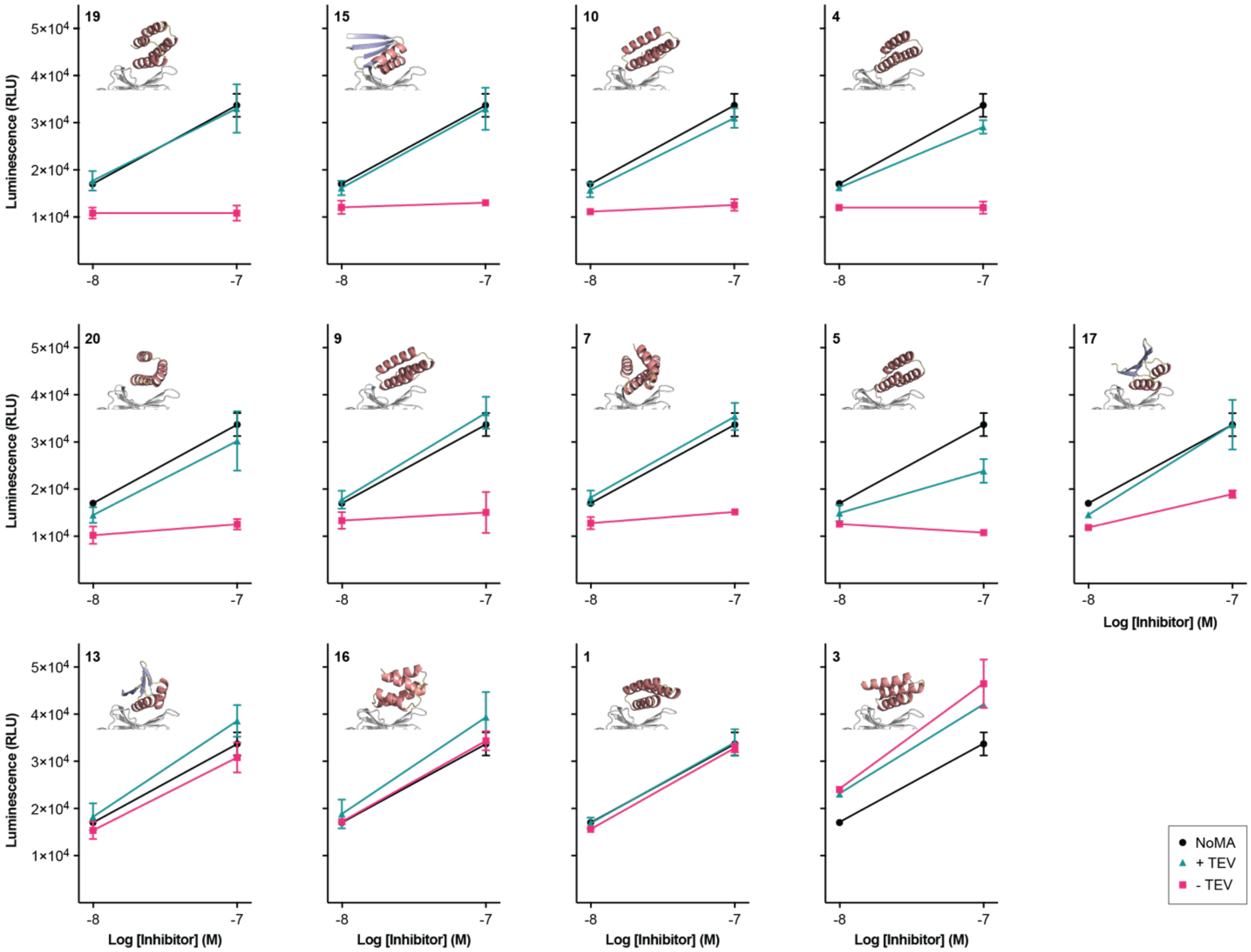


**Supplemental Figure 3 Masked antagonist cellular activity screen.**
In the assay, when the endogenous PD-1:PD-L1 interaction is disrupted, TCR activation induces luminescence. To compare the activity of the various masked inhibitors at 10 and 100 nM, the luminescence responses without (pink) and with (teal) TEV incubation are shown. To directly compare, the activity of NoMA (black) is also shown in each panel. Two technical replicates from a single experiment, presented as mean ± SEM, are shown. Across the figure from left to right, each masked inhibitor is shown in order of the largest to smallest difference upon protease treatment at 100 nM. A cartoon representation of the masked antagonist model is also shown (HA-PD1 in grey and the AiD colored by secondary structure).

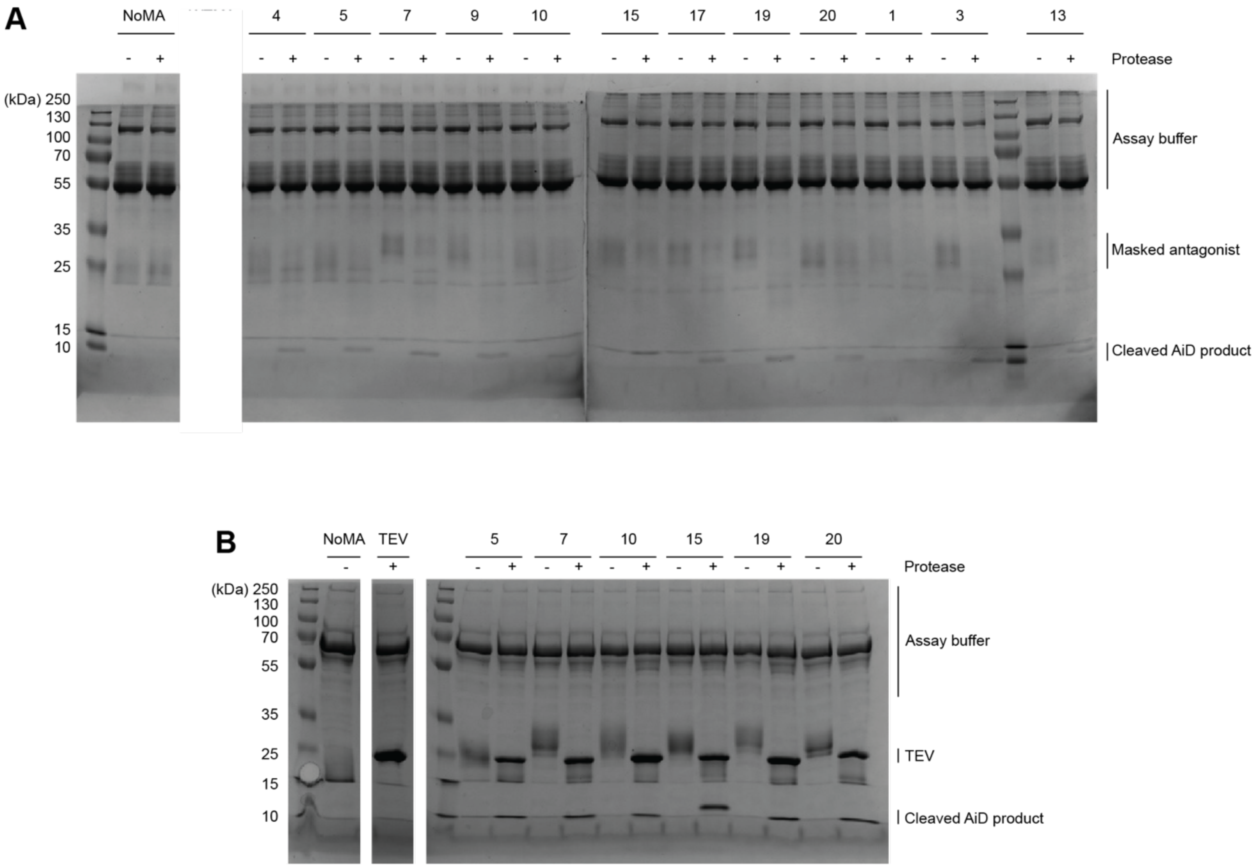


**Supplemental Figure 4. Cleavage of masked antagonists during cellular assay.** Prior to the Promega cellular blockade assay screen **(A)** or the full assay **(B)**, masked antagonists were incubated with and without protease overnight. After the assay, representative samples were run on an SDS-PAGE (16% acrylamide) gel. As positive controls, NoMA was cleaved in the same manner. Smearing is due to glycosylation from the Expi293F mammalian expression system.

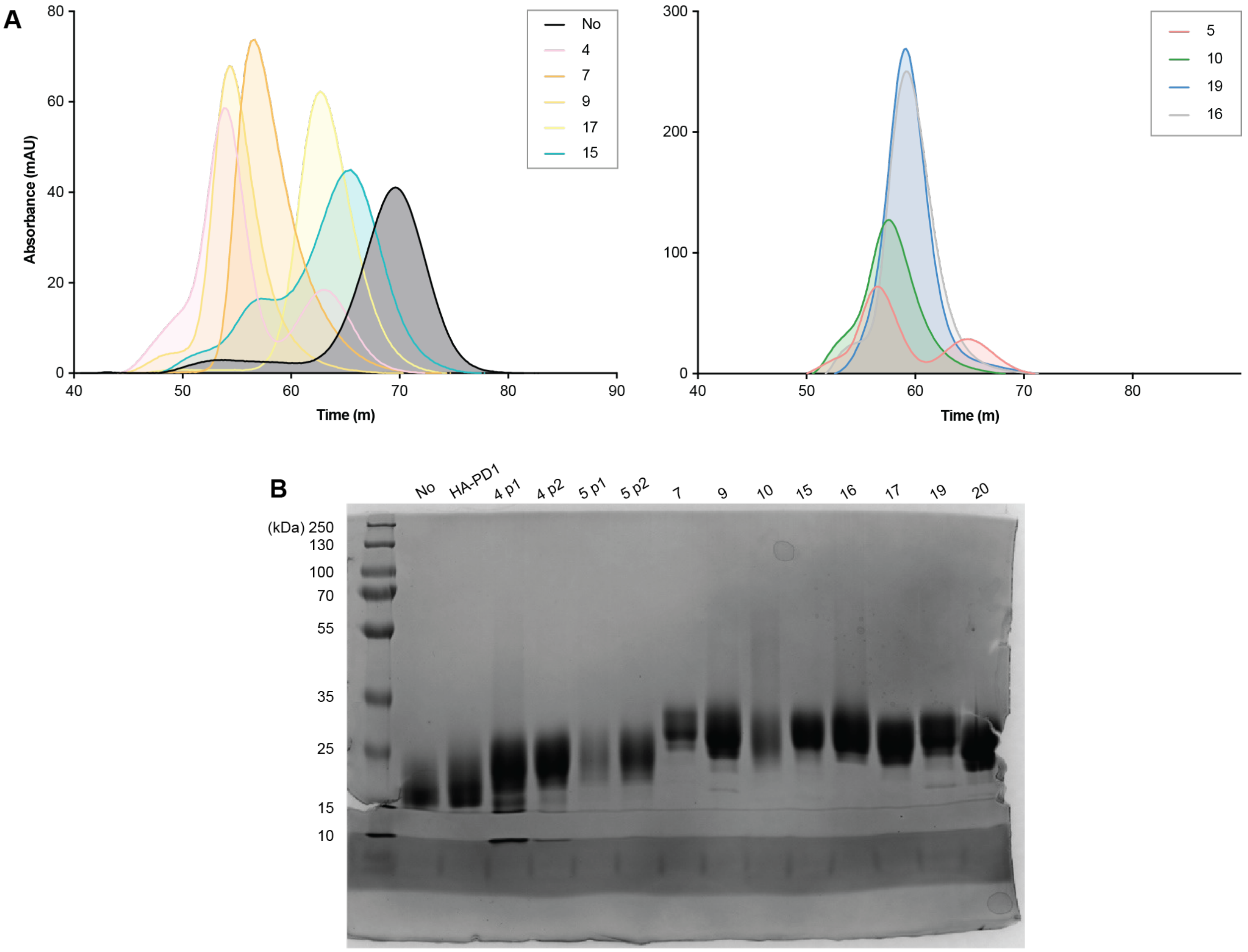


**Supplemental Figure 5. Expression and purification of masked antagonists.**
**(A)** Size exclusion chromatography (HiLoad Superdex 75 pg) traces of the 10 masked antagonists after expression in a 125 mL flask and then followed by nickel affinity chromatography. As MA5, MA10, MA16, and MA19 were purified on a different HiLoad Superdex 75 pg column, they are shown separately (right). **(B)** SDS-PAGE (16% acrylamide) of the 10 masked antagonists. For MA4 and MA5, both peaks were run separately. As positive controls, NoMA ("No") and HA-PD1 were purified in the same manner. Smearing is due to glycosylation from the Expi293F mammalian expression system.

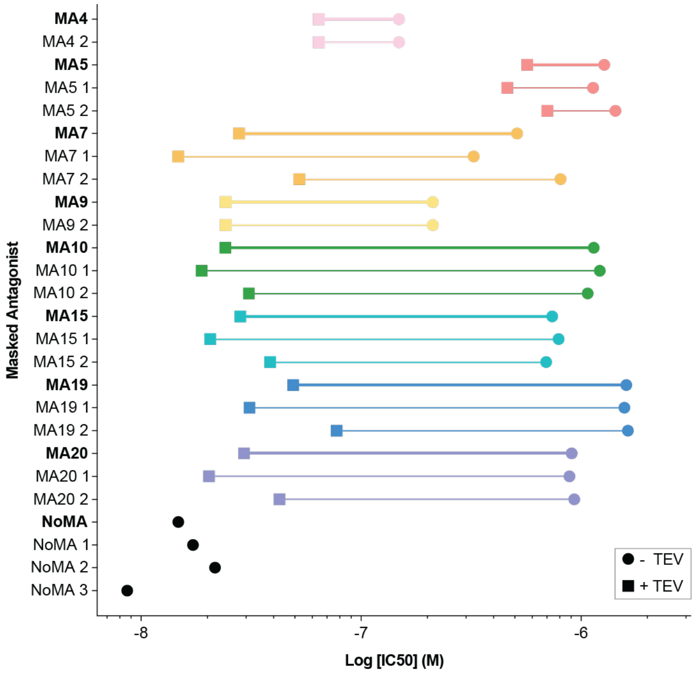


**Supplemental Figure 6. Masked antagonists show protease-dependent PD-L1 binding.**
In the cell surface blockade assay, when the endogenous PD-1:PD-L1 interaction is disrupted, TCR activation induces luminescence. Data from three separate experiments were analyzed in Prism and the mean LogIC50 windows with (square) and without (circle) protease incubation for each masked PD-L1 antagonist are shown. The average (bold) and individual values from the three separate experiments (1, 2, and 3) are shown.

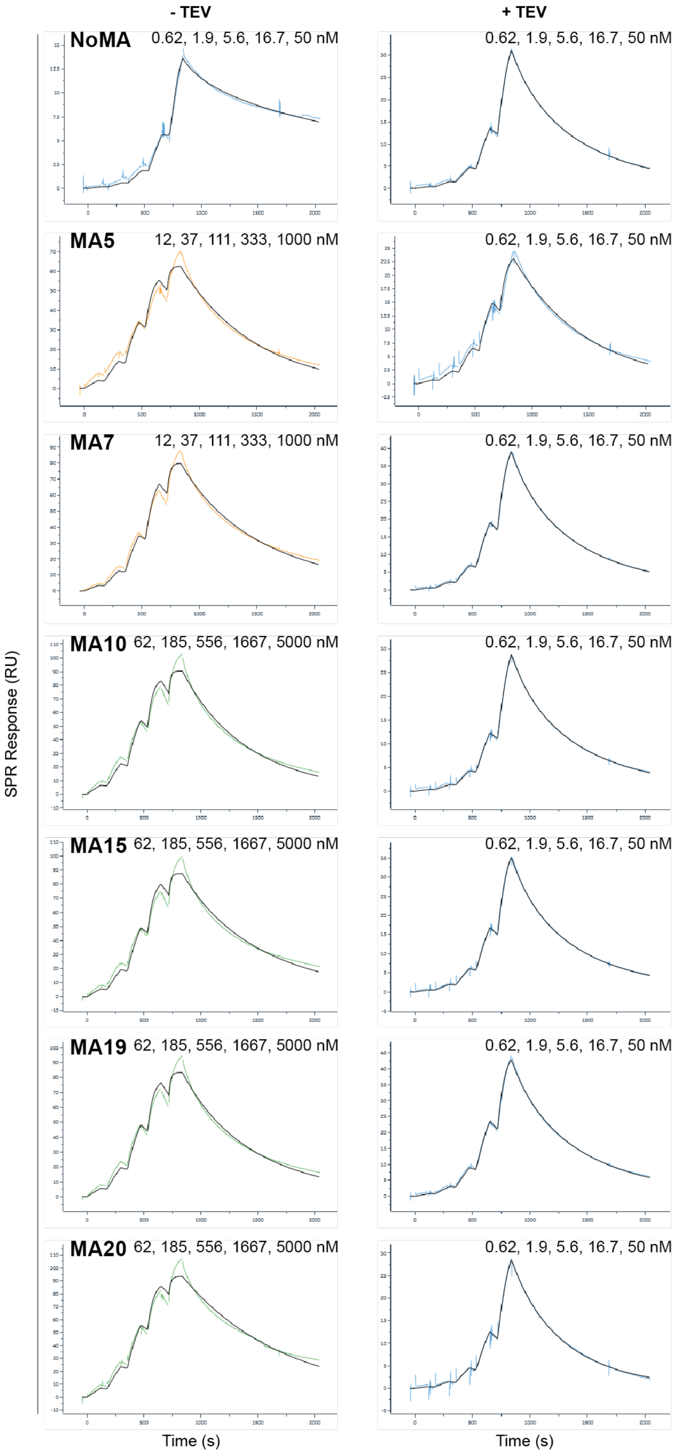


**Supplemental Figure 7. Representative SPR sensorgrams of the various masked antagonists at various concentration ranges.**
Representative SPR sensorgrams (colored lines) and fits (black) as generated by the Biacore Insight Evaluation software, for the binding kinetics of masked antagonists with and without protease treatment to biotinylated PD-L1.

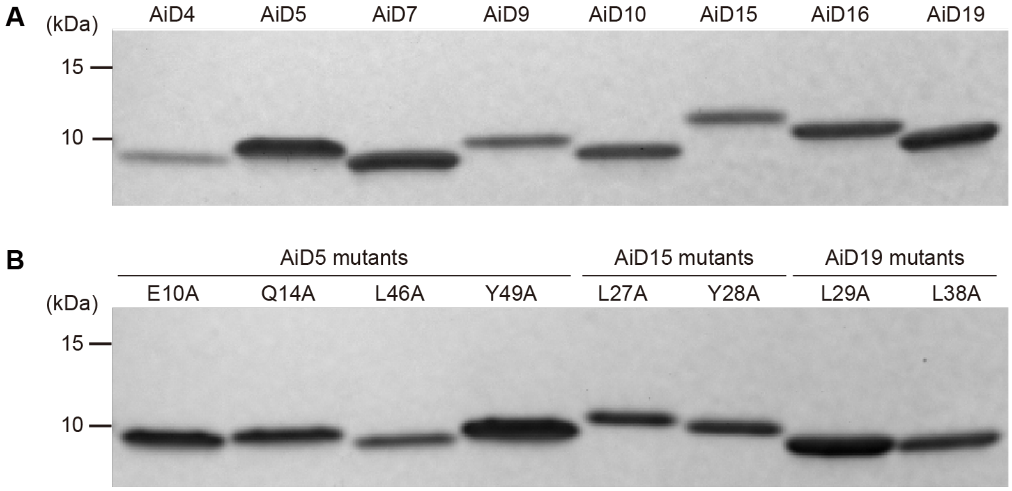


**Supplemental Figure 8. Expression and purification of recombinant AiDs and mutants.**
SDS-PAGE (4-20% acrylamide) analysis of various AiDs **(A)** and AiD mutants **(B)** obtained with Ni-affinity purification followed by gel filtration. The gel was stained with Coomassie Brilliant Blue.

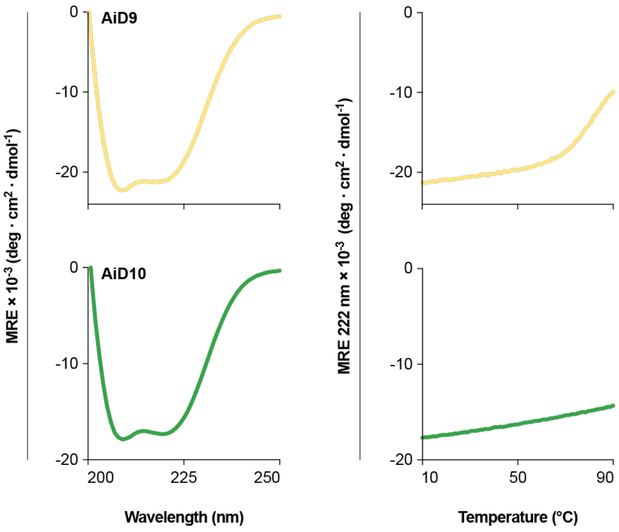


**Supplemental Figure 9. Additional CD spectra of recombinant AiDs.**
Additional CD spectra of recombinant AiD9 and AiD10 display negative peaks at 208 and 222 nm, consistent with the formation of α-helices (left) and temperature melts demonstrated thermostability (right).

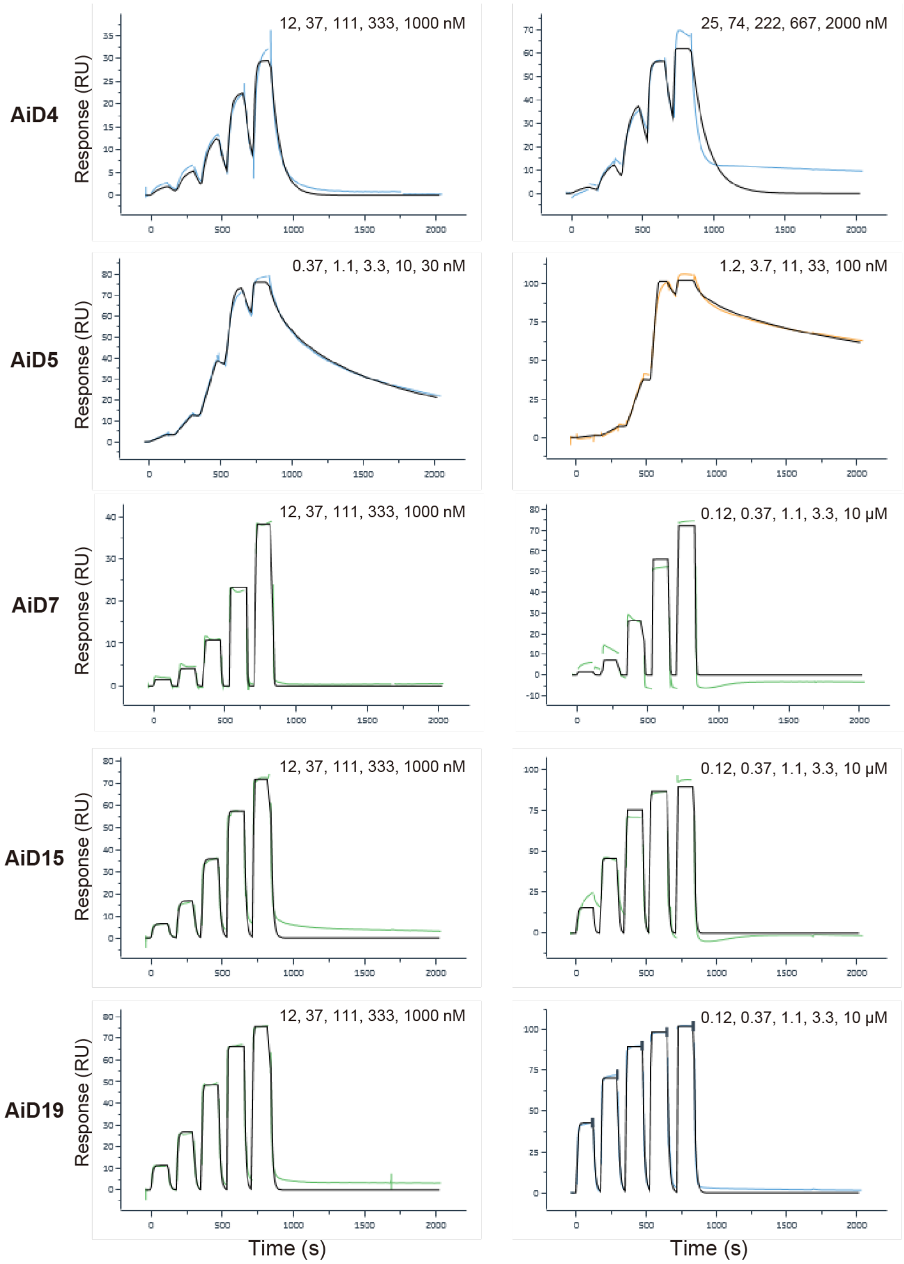


**Supplemental Figure 10. Representative SPR sensorgrams of the various recombinant AiDs at various concentration ranges.**
Additional representative SPR sensorgrams (colored lines) and fits (black) as generated by the Biacore Insight Evaluation software, for the binding kinetics of recombinant AiD proteins to biotinylated HA-PD1.

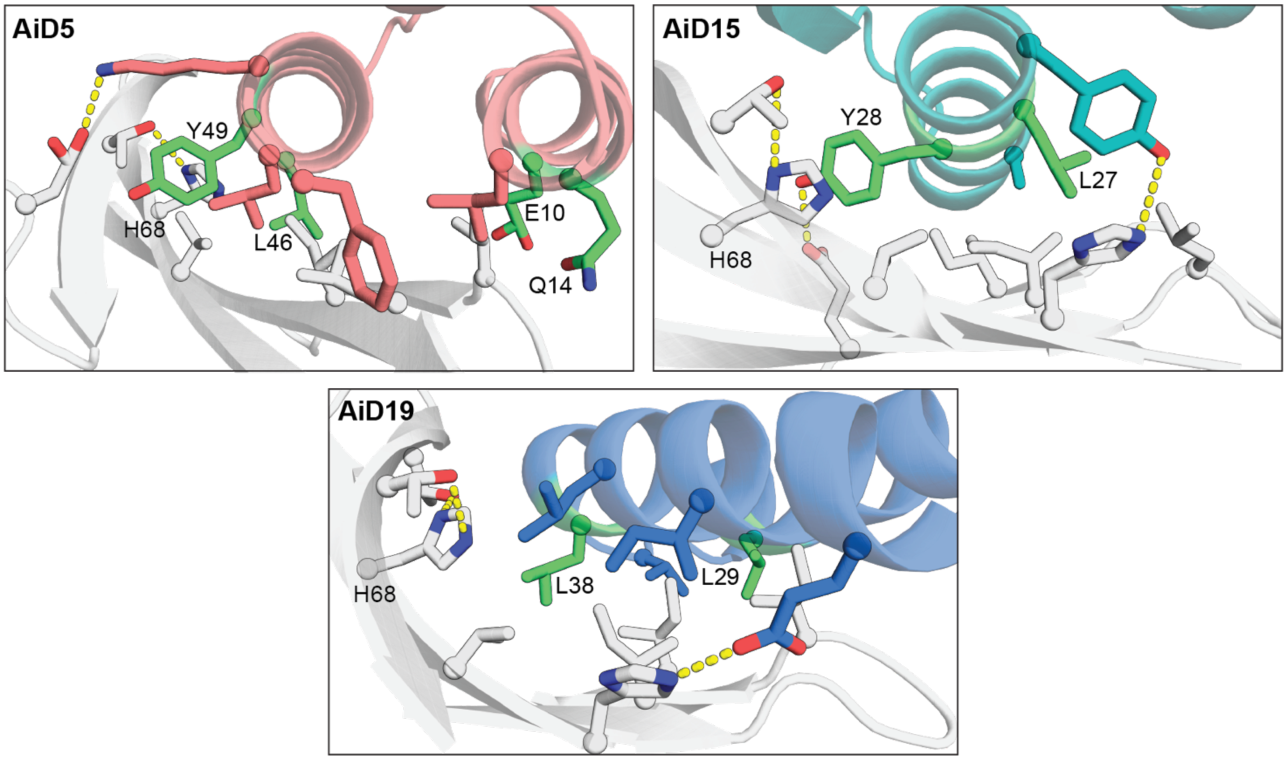


**Supplemental Figure 11. Mutation of key interface residues to disrupt binding.** Interface residues for single alanine mutation constructs (green) for binding studies for the design models AiD5, AiD15, and AiD19. Figure created using PyMOL.

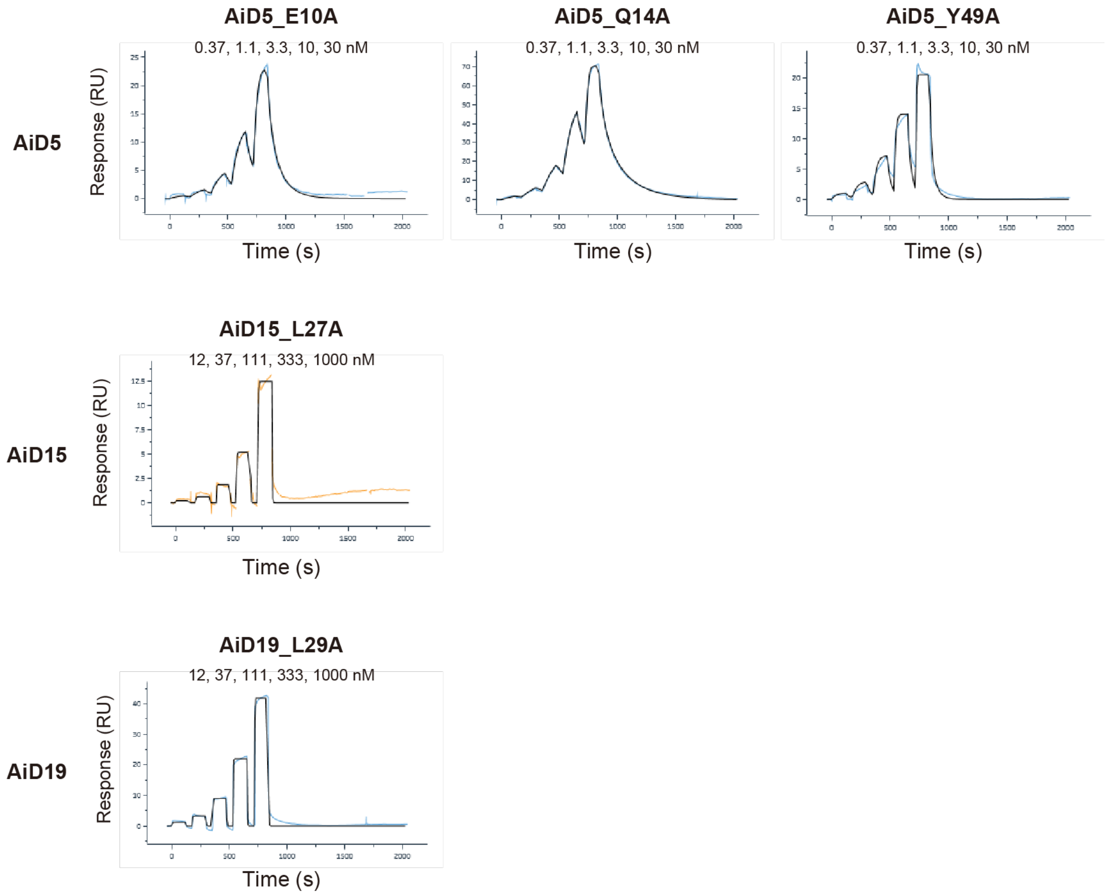


**Supplemental Figure 12. Representative SPR sensorgrams of the mutant recombinant AiDs at various concentration ranges.** Additional representative SPR sensorgrams (colored lines) and fits (black) as generated by the Biacore Insight Evaluation software, for the binding kinetics of recombinant AiD mutants to biotinylated HA-PD1.
